# Species interactions and diversity: a unified framework using Hill numbers

**DOI:** 10.1101/2022.10.18.512607

**Authors:** William Godsoe, Rua Murray, Ryosuke Iritani

**Affiliations:** Department of Pest Management and Conservation, Lincoln University, PO Box 85084, Lincoln 7647, New Zealand; School of Mathematics and Statistics, Univ. of Canterbury, Private Bag 4800, Christchurch, 8140 New Zealand; RIKEN Interdisciplinary Theoretical and Mathematical Sciences (iTHEMS), Wako, Japan

**Keywords:** Biodiversity, metacommunity, Hill number, competition, mutualism, predation, partition, relative fitness

## Abstract

Biodiversity describes the variety of organisms on planet earth. Ecologists have long hoped for a synthesis between analyses of biodiversity and analyses of biotic interactions among species, such as predation, competition, and mutualism. However, it is often unclear how to connect details of these interactions with complex modern analyses of biodiversity. To resolve this gap, we propose a unification of models of biotic interactions and measurements of diversity. We show that analyses of biodiversity obscure details about biotic interactions. For example, identical changes in biodiversity can arise from predation, competition or mutualism. Our approach indicates that traditional models of community assembly miss key facets of diversity change. Instead, we suggest that analyses of diversity change should focus on partitions, which measure mechanisms that directly shape changes in diversity, notably species level selection and immigration, rather than traditional analyses of biotic interactions.

**Speculations:** Our paper proposes that observations of biodiversity cannot be used to distinguish different types of biotic interactions. For generations ecologists have been fascinated with the links between biodiversity and biotic interactions (i.e. competition, mutualism and predation). Many of us expect that observations of biodiversity provide vital clues about how biotic interactions operate in nature, but it is hard to tell when these clues are reliable.

Our work integrates models of biotic interactions and measurements of biodiversity diversity change. This highlights how an observed change in diversity can be compatible with any type of biotic interaction (i.e. competition, mutualism, predation etc.). So for example, the same increase in Shannon diversity could indicate the superior ability of a competitor the success of mutualists or a predator’s tendency to harvest dominant prey species. This is so because diversity measures are designed to be concerned with changes in relative abundances but not account for absolute abundance changes.

Observations of biodiversity change are unlikely to yield insights about biotic interactions per se because biodiversity itself obscures species’ absolute abundances. Therefore, models of diversity change should focus on mechanisms that are less influenced by changes in absolute abundances such as species-level selection.

## Introduction

Ecologists have long sought to link dynamic models of biotic interactions to better understand changes in biodiversity (Chesson 2000; MacArthur 1965; MacArthur 1972; Urban et al. 2016; Vellend 2016). Examples of biotic interactions include competition, mutualism and predation. Biodiversity describes the variety of living things, for example species richness. Overall measurements of diversity (gamma diversity) are commonly partitioned into diversity within locations (alpha diversity) and dissimilarities among locations (beta diversity) (Jost 2006; Jost 2007). One of the most common sources of evidence for the effects of species interactions is spatial patterns in diversity. In a world facing extensive anthropogenic disturbance, we need to understand the mechanisms that shape biodiversity. To do this, we propose a unification of dynamic models of biotic interactions with measurements of diversity.

Observations of diversity have long been used as a tool to understand the role of species interactions (Hubbell 2001; MacArthur 1965; Palmer 1994; Sax et al. 2002; Simha et al. 2022). For example, Chomicki et al. (2019) proposes a suite of mechanisms through which mutualists shape community assembly, the most common source of evidence is observed patterns of diversity. Generations of community ecologists have used observations of species high diversity of species in nature as evidence for competition mediated by niche differentiation among species (Simha et al. 2022). However, it is unclear when observations of diversity can be used to tease apart the effects of biotic interactions (Barner et al. 2018; Poisot et al. 2015; Vellend 2010).

What has been missing is a common framework for models of biotic interactions and models of species diversity. This gap is exemplified in Robert H. MacArthur’s classic review of patterns of species diversity that includes a theory section devoted to dynamic on models of resource competition, across spatial scales (MacArthur 1965). Unfortunately, MacArthur presents a separate model for pattern (biodiversity) and process (competition), each with different units, formulae and graphical analyses. Similarly Hill 1973 states that diversity is of theoretical interest because it can be related to attributes of communities such as predation pressure (a species interaction) and stability (an outcome of species interactions). Both Hill and MacArthur envision links between species interactions and diversity, but neither shows what this formal connection should look like. This omission suggests the need for further synthesis, where the connections between analyses of diversity and analyses of biotic interactions are placed in a common framework. Such clarifications can help us to understand what is missing when ideas from one framework are translated to another.

Subsequent work has emphasized simplified models of biotic interactions, such as the neutral theory of biodiversity (Hubbell 2001), where all interactions are assumed to be competitive and all individuals of any species are assumed to be similar. Similar models, with similar limitations have been proposed by evolutionary biologists to study the diversity of genotypes within a species (Lewis et al. 2018; Sherwin et al. 2017), or modern coexistence theory where interactions are studied close to equilibrium (Barabás et al. 2018; Chesson 2000). It is now clear that assumptions such as zero sum dynamics close to equilibrium are unlikely to hold in nature (Callaway et al. 2002; Fukami and Nakajima 2011). The time is right to relax this “zero-sum” assumption and explicitly consider what information about biotic interactions can be extracted from analyses of biodiversity.

One impediment to linking biotic interactions and biodiversity has been the ambiguity in how biotic interactions are defined (Abrams 1987; Hart et al. 2018; Rees et al. 2012). To mitigate this ambiguity we will define biotic interactions based on their effects on absolute abundances (Abrams 1987; Chamberlain et al. 2014; Holland and DeAngelis 2010). This means that competition occurs when species reduce each other’s absolute abundances. Mutualism occurs when species increase each other’s absolute abundances and predators reduce the absolute abundances of their prey.

It has long been hoped that observational studies can distinguish different types of biotic interactions. Notably, some analyses have suggested that competitive interactions stabilize communities while mutualistic interactions destabilize communities (Stone 2020) and predatory interactions frequently produce oscillations (Holt 2011). Despite the likely implications for diversity, these links are difficult to analyse using current methods (Urban et al. 2016).

The causes of diversity change in nature can be difficult to tease apart. Over long time scales, biotic interactions shape biodiversity, such as when introduced predators lead to the extinction of island endemics (Bellingham et al. 2010), or low elevation plant species outcompete alpine species (Alexander et al. 2015). However, it is often difficult to distinguish the effects of biotic interactions from a suite of mechanisms including physiological effects, dispersal, and the abiotic environment (Germain et al. 2018; Urban et al. 2016). Biotic interactions are also prone to change across environmental gradients (Chamberlain et al. 2014; Louthan et al. 2015); for example, interactions among many plant species are competitive in benign environments but shift to mutualistic in stressful environments (Callaway et al. 2002; Chamberlain et al. 2014). In some communities *Greya* moths acts as a mutualist to the wildflower *Lithophragma parviflorum* because of the benefits it provides from pollination whereas in other communities the same moth acts as an antagonistic seed predator (Thompson and Cunningham 2002).

Different biotic interactions can produce similar effects on biodiversity. For example, analyses of long-term species coexistence generalize to other interactions including mutualisms and predation (Lanuza et al. 2018; Spaak et al. 2021; Venail et al. 2014). Likewise competition for resources can be indistinguishable from a radically different mechanism, indirect effects of predation (Chesson and Kuang 2008; Holt 1977). Many short-term effects of biotic interactions are also comparable. For example laboratory experiments with fruit flies (*Drosophila* spp.) show that negative frequency dependence can arise among competitors (Ayala et al. 1973), meaning that each competing species has an advantage when it is rare. Similar patterns of frequency dependence have been observed between predators and prey (Kihara et al. 2011), and among mutualists (Harcombe et al. 2018). Rapid changes in biodiversity may be confounded by additional complications such as extinction debt (Catford et al. 2018; Gilbert and Levine 2013) tipping points and chaos (Chase et al. 2019; Fukami and Nakajima 2011; Hastings 2004; Storch et al. 2021). These similarities among biotic interactions have led to the development of a common theoretical framework for describing population dynamics (Godsoe et al. 2017a; Holland and DeAngelis 2009; Holland and DeAngelis 2010), but the implications for analyses of diversity remain unclear.

There are many different ways to measure diversity (Magurran and McGill 2011), making it prohibitively difficult to link every possible measurement to biotic interactions. Fortunately, many measurements of biodiversity can be organized into a single, analytical framework with common units known as Hill numbers (Hill 1973; Jost 2006; Jost 2007). This framework encompasses analogues to traditional diversity measurements within a community including species richness, Shannon entropy and Simpson’s diversity. It also includes methods to partition overall diversity in a metacommunity (gamma diversity) into average diversity within communities (alpha diversity) and diversity due to differences between communities (beta diversity) (Chao and Chiu 2016). This framework makes it possible to synthesize changes in diversity across measurements and across scales. Hill numbers thus make a convenient tool to link of previous analyses of diversity and biotic interactions.

In view of the difficulties of linking biotic interactions and biodiversity, some have proposed analyses of diversity change that do not explicit model biotic interactions. For example, Vellend (2016) focuses on models of selection among species a mechanism, which implicitly captures effects of biotic interactions. Vellend’s framework emphasizes conceptual insights, but several authors have proposed empirical approaches inspired by this insight. For example, Godsoe et al. (2022) showed how models of selection could be used to analyse changes in Shannon diversity at multiple spatial scales, while Tatsumi et al. (2021)analysed changes in measures of diversity related to species richness. Though these approaches show promise we currently lack the ability to explicitly compare these explicit and implicit models of biotic interactions.

Here we propose a unification of biotic interactions and diversity measurements using techniques from evolutionary theory (Lion 2018). The goal of this unification is to clarify the connections between mathematical models of biotic interactions and diversity. Using the replicator equation from evolutionary theory as a unifying concept. This connection makes clear that different types of biotic interactions produce indistinguishable changes in diversity. We illustrate this first through graphical models then for a general model of change in a metacommunity. Our framework suggests three key results 1) measurements of diversity obscure changes in absolute abundances 2) as a result changes in diversity often cannot be used to distinguish biotic interactions. 3) Context dependency across a metacommunity limits our ability to distinguish biotic interactions with diversity change. In view of these limitations, we propose that better predictions can be made by linking shifts in species frequencies to shifts in diversity.

## The model

In this section, we use a graphical model to illustrate how diversity obscures the effects of biotic interactions. To do this, we consider two species with absolute abundances *n*_1_(*t*) and *n*_2_(*t*) in a single community (at a given continuous-time point *t* unless otherwise stated). We assume that these numbers are known with certainty so that there are no artefacts due to sampling.

One way to define biotic interactions uses the effects of one species on the absolute abundances of another (Godsoe et al. 2017b; Holland and DeAngelis 2009; Holland and DeAngelis 2010; Odum 1983). Using this approach, an interaction is competitive when it causes a decline in the absolute abundances of both species, relative to their growth rate in the absence of the interaction. For example the fruitfly species *Drosophila willistoni and D. pseudoobscura* cause declines in each other’s abundances when they co-occur (Ayala et al. 1973). Interactions that are mutualistic cause an increase in the absolute abundances of each species, such as when different plant species increase each other’s abundances by mitigating the effects of stressful environments (Callaway et al. 2002). We treat facilitation as a form of mutualism (Chamberlain et al. 2014). A predator will cause a decline in the absolute abundance of prey and prey cause an increase in the abundance of predators. These effects of biotic interactions can be illustrated in a plot of the absolute abundance of species 2, versus species 1 (Fig. 1 A; See Appendix S2 for R scripts used to generate figures).

**Figure 1.**
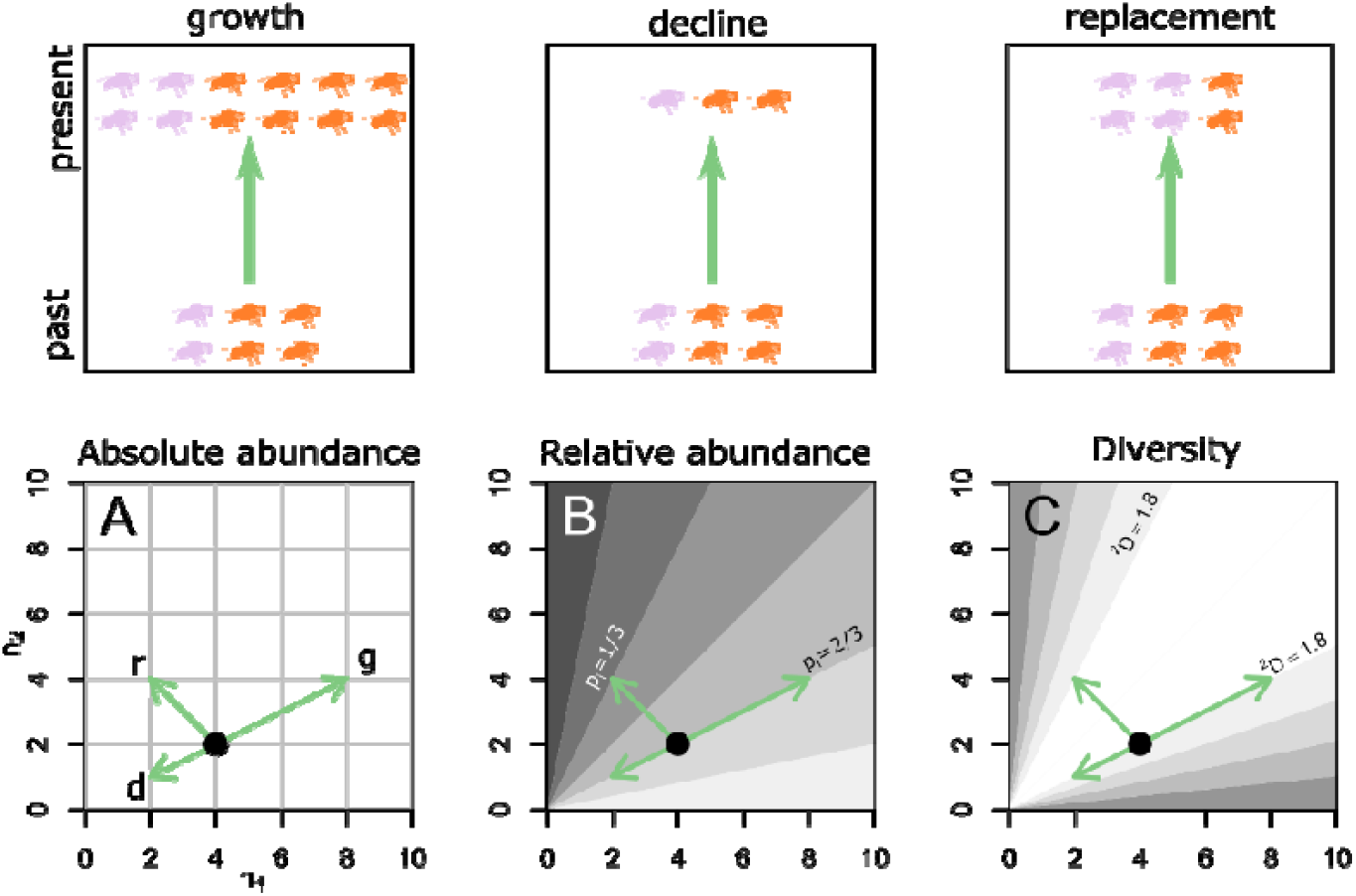
Different changes in absolute and relative abundances are not necessarily reflected in the change in diversity. The top row shows three ways absolute abundance can change, growth in all species (g), decline in all species (d), replacement of one species by another (r). A) All three scenarios change the absolute abundance of species 1 (*n*_1_) and (*n*_2_), each scenario is depicted as a green arrow and labelled growth (g), decline (d) and replacement (r). B) Shades the same plot by relative abundance of species 1 (n_1_) going from low relative abundances (dark grey) to high relative abundances (light grey). Using relative abundances makes the growth and decline scenarios indistinguishable. C) Shades the plot by diversity (the Hill number equivalent of Simpson’s ^*2*^*D*). Using diversity all three scenarios are indistinguishable, with each representing no change.

Biodiversity does not depend directly on absolute abundances, it depends instead on relative abundances. The relative abundance of species 1 is 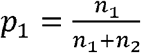, where *n*_1_, *n*_2_ is the number of individuals of species 1 and 2 respectively. The relative abundance of species 2 is 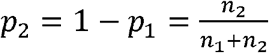. The same absolute abundances can imply very different relative abundances (Fig. 1B). For example a community the relative abundance of species 1 is 2/3, may be a community where with 2 of 3 individuals belonging to species 1 or a community where 4 of 6 individuals belong to species 1, or a community where 8 of 12 individuals belong to species 1 or many other possibilities.

Measurements of diversity further obscure information on abundances. This is because diversity ignores information on species identity. For example, diversity a community with two flies of species 1 and one fly of species 2 has the same diversity as a community with one fly of species 1 and two flies of species 2 (Fig. 1 C). Fig. 1 uses one diversity measurement, the Hill number associated with Simpson’s diversity, where the ambiguity is illustrated by the fact that the graph is diagonally symmetrical meaning that diversity is equivalent for pairs of values on either side. This symmetry is shared by all measures of species diversity in a community (Leinster 2021). Appendix S1 Fig. S1 illustrates equivalent plots for other diversity measurements.

Fig. 1 illustrates three distinct changes in absolute abundances, population growth, population declines and the replacement of one species by another. Each of these changes in absolute abundances (Fig. 1 A) fail to change diversity (Fig. 1 C). This figure captures the central message of our manuscript: diversity can obscure changes caused by biotic interactions. We will formalize this claim in our analysis of Equation 4 below. With more species, the geometry is more difficult to visualise, and so in the next section, we extend this framework to multiple species using an analysis of dynamic models of biotic interactions and biodiversity.

### Dynamics of biotic interactions

To model biotic interactions, we consider a metacommunity which can be divided into species labelled *i*=1,2,….*I* which inhabit communities *j*=1,2,…,*J*. The number of individuals of species *i* in community *j* is denoted *n*_*i,j*_, hereafter deme (*i,j*). Changes in absolute abundances for each deme are given by:

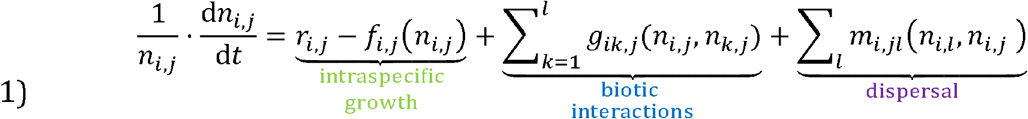

This represents the per-capita population growth rate of species *i* in community *j*. This growth rate is a function of species *i*’s intrinsic growth rate *r*_*i,j*_, density dependent interactions among individuals of species *i, f*_*i,j*_(*n*_*i,j*_) and interactions between species *i* and an other species *k, g*_*ik,j*_. Competition occurs when each species *i* reduces the growth rate of species k and vice versa (*g*_*ik,j*_, *g*_*ki,j*_ <0). Mutualism occurs when each species increases the growth rate of the other (*g*_*ik,j*_, *g*_*ki,j*_ >0). A consequence of predation is that one species (the prey) increases the growth rate of the other (the predator). Interactions within and among species may be non-linear (Holland and DeAngelis 2009).

The dispersal term describes movement into deme (*i,j*) from other demes (which we index by *l*). The expected number of individuals of species *i* dispersing to community *j* from community *l*, per unit time is *m*_*i,jl*_(*n*_*i,l*_, *n*_*i,j*_)*n*_*i,j*_. All individuals disperse within the metacommunity such that ∑_*j*_ *m*_*i,jl*_ (*n*_*i,l*_,*n*_*i,j*_)*n*_*i,j*_ = 0. This approach is based on Gravel et al. (2016), and allows for any spatial arrangement of communities (i.e. communities may be organized in a lattice or network). See Appendix S1 for an example of how to parameterize the interesting special case of passive dispersal in our framework.

Changes in absolute abundances described in Equation 1 can be converted into changes in relative abundances. This is done by changing the variable of interest to the relative abundance of deme 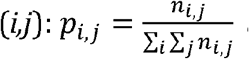, and the total number of indviduals across the metacommunity *N* = ∑_*i*_ ∑_*j*_ *n*_*i,j*_. This leads to the dynamic equations (Hofbauer and Sigmund 1990; Lion 2018; Taylor and Jonker 1978):

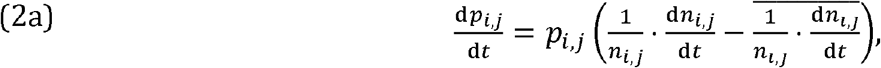

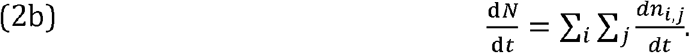

Equation 2a is commonly known as the replicator equation (Lion 2018). It is one way to describe the relative fitness of each deme. Equation 2a states that the proportion of the metacommunity belonging to species *i* in community *j* will increase when that deme has a higher per-capita growth rate 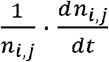, then the average per-capita growth rate across the metacommunity 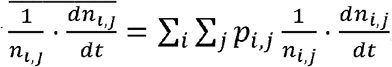. Changes in the total number of individuals are captured by Equation 2b. In effect moving from Equation 1 to Equation 2a is like moving from Fig. 1 A, that tracks changes in absolute abundances of all species to Fig. 1b that tracks changes in relative abundances. Though the replicator equation is well known in evolutionary theory (Nowak 2006), it has been less clear how to relate this equation for change in relative abundances into change in contemporary measurements of diversity. In the next section we will derive a new expression linking the replicator equation and diversity change (equation 4).

Many individual metrics of diversity can be organized into a common framework (Gaggiotti et al. 2018; Hill 1973; Jost 2006; Jost 2007; Sherwin et al. 2017)—i.e., Hill numbers. All Hill numbers are summaries of information on the relative abundance of demes. For example, gamma diversity summarizes information on each species’ relative abundance across the metacommunity ignoring spatial subdivision:

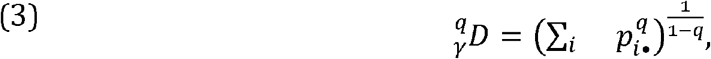

The relative abundance of species *i* is simply the sum of the relative abundances of species *i* across all demes (*p*_*i•*_ = ∑_*j*_ *p*_*i,j*_). The parameter *q*, hereafter the diversity order changes the emphasis placed on species at low relative abundance; *q* can represent any real number. Low diversity order produces a diversity index that emphasizes species at low relative abundance. For example, species richness is recovered when *q*=0. Higher diversity orders increase the emphasis on common species such as *q*=2 Simpson’s diversity. In the limit that *q* approaches 1 Equation 4 approaches Shannon Wiener diversity, the exponential of Shannon entropy (Jost 2006; Jost 2007).

We can derive a general expression for change in gamma diversity by computing the derivative of Equation 3 with respect to time using the chain rule.

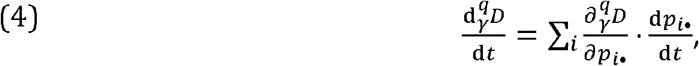

In words, Equation 4 represents change in diversity by computing the change in species’ frequencies with respect to time 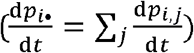, then translating changes in species’ frequencies into changes in diversity 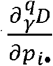, (note the *∂* denotes partial derivatives). We provide examples of this translation in Fig. 2. There is no exact analogue to this term in traditional analyses of community assembly (Adler et al. 2007; Chesson 2000; Hubbell 2001; Vellend 2016). Equation 4’s function is analogous to the move for analyses of relative abundances in Fig. 1b to analyses of diversity in Fig. 1c, but Equation 4 is applicable to any number of species.

**Figure 2.**
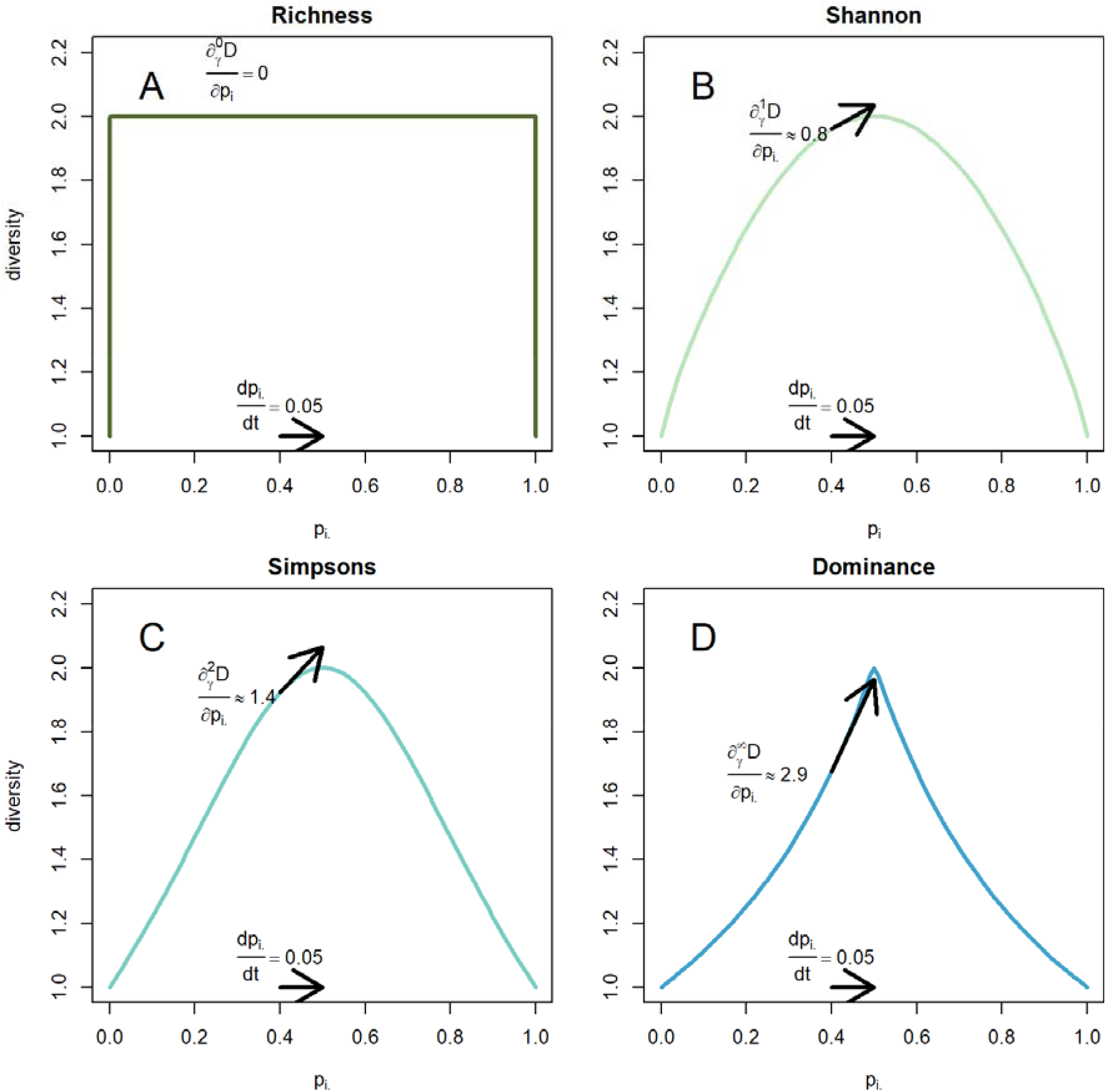
Illustrates of how changes in relative abundance translate into changes in diversity. Each pane is a plot of diversity versus the relative abundance of one species (*p* _*i*•_, in a two species metacommunity). Each panel shows the consequence of an increase in the relative abundance of a moderately common species 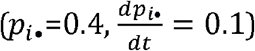. A) This results in no change in richness 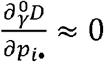, B) a small increase in Shannon 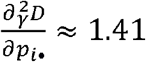, C) a larger increase in in Simpsons’ diversity and D) a still larger increase in Dominance 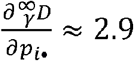.

Interactions that are relatively easy to distinguish using information on absolute abundances can lead to surprisingly similar effects on diversity. Fig. 3 illustrates four sets of simulations for population dynamics of two species, each starting at the same initial conditions. As an aid we have also included absolute abundance each species reaches in the absence of the other, i.e. the carrying capacity of species 1 (purple vertical line) and species 2 (orange purple line). In the absence of species interactions (a state sometimes known as neutralism), populations of both species grow until they reach their carrying capacity (Fig. 3 A). In the face of competition, the absolute abundances of both species decline below their carrying capacity (Fig. 3 C). In the face of mutualism each species’ absolute abundances exceed their carrying capacity (Fig. 3 E) and in the face of predation the prey (species 1) declines below its carrying capacity while the predator (species 2) exceeds its carrying capacity (Fig. 3 F). All examples start at the same initial conditions, and arrive at a final diversity of ^*2*^*D*=1.8. Each simulation produce qualitatively similar rises and falls in diversity over time (Fig. 3 B, D, E, G).

**Figure 3.**
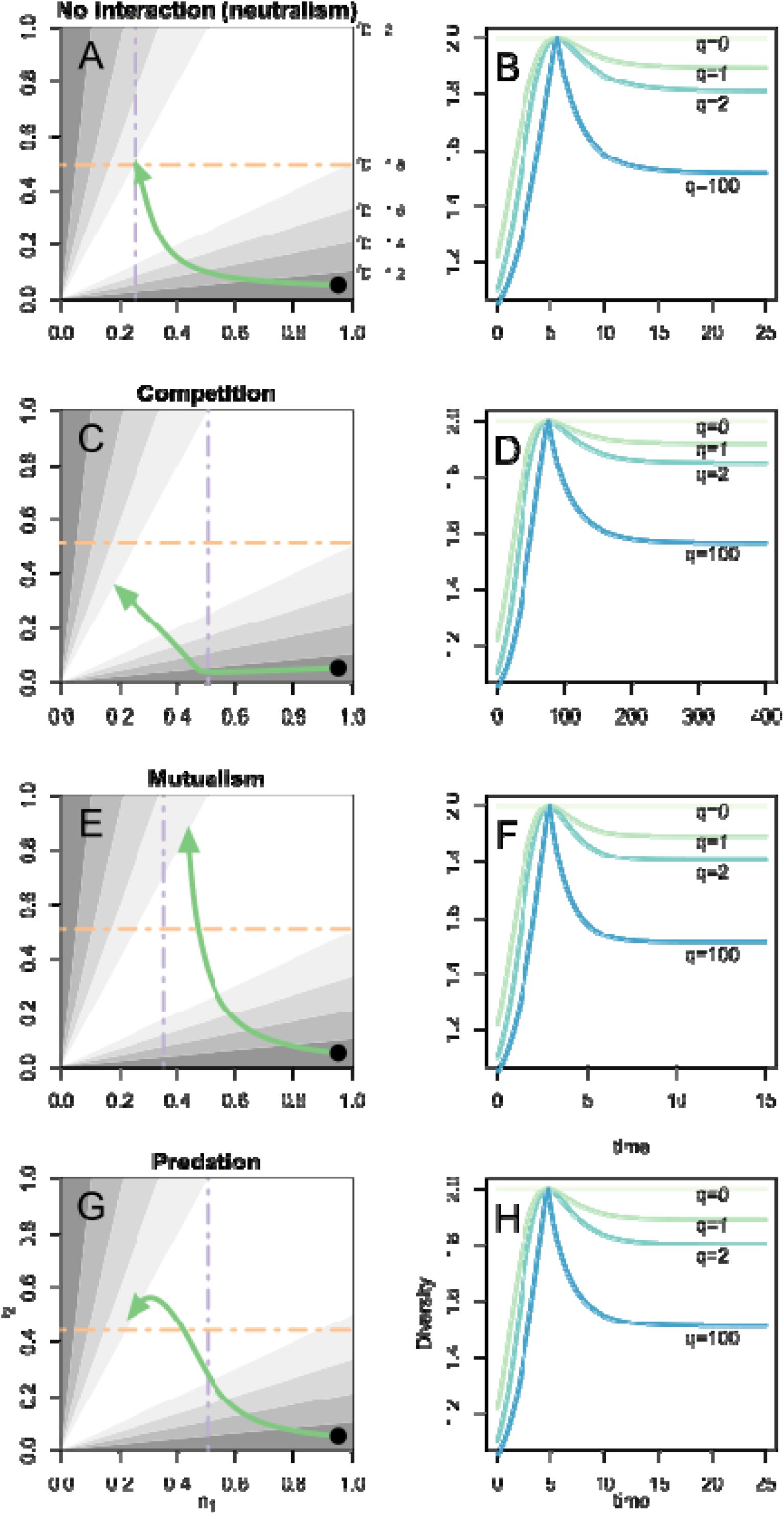
Illustrates how different biotic interactions have similar long-term effects on diversity. A) Illustrates changes in absolute abundances of species 1 and species 2 (green line) in a simulation of neutralism (i.e. the absence of biotic interactions). In this simulation each species ultimately reaches its carrying capacity (the point where the yellow and purple lines cross). Shading depicts the numbers equivalent of Simpson’s diversity for each combination of absolute abundances. B) This leads to a rise and fall in most Hill numbers as species 2 goes from rare to common (increase in diversity) then becomes more common than species 1 (decrease in diversity). Species richness, remains unchanged 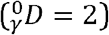. C) Simulates the effects of competition on the absolute abundance. The absolute abundances of species 1 and 2 decline below their carrying capacities. D) Over the course of the simulation most diversity orders increase then decline. E) Mutualism increases the absolute abundances of both species above their respective carrying capacities. F) This leads to comparable changes in diversity, as species 2 increases in relative abundances. G) Predation decreases the absolute abundance of species 1 (the prey) and increases the absolute abundances of species 2 (the predator). H) This leads to comparable changes in diversity. Simulations based on a Lotka-Volterra model parameter values (Appendix S1 table S1).

### Diversity change across a metacommunity

The difficulties in identifying biotic interactions remain when diversity is studied across a metacommunity. When multiple communities are considered, gamma diversity is partitioned into a measure of average local diversity (hereafter alpha diversity) and a measure of community dissimilarity (hereafter beta diversity). For most diversity measurements, Jost (2007) recommends defining alpha diversity as the diversity within each community, averaged across all communities:

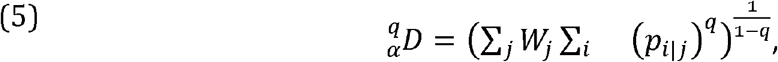

where *p*_*i*|*j*_ represents the relative abundance of species *i* in community 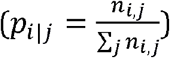. The contribution of each community to overall diversity is assigned a weight *w*_*j*_. Jost (2007) proposes a general expression for these weights 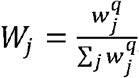, but recommends that all communities be weighted equally, for most diversity measurements *w*_*j*_ = 1/*J*. However, analyses of biotic interactions across a metacommunity tend to weight all individuals equally (Chesson et al. 2005). Only one measurement is suitable for weighting individuals across the metacommunity equally; Shannon diversity, which is found when *q*=1.

Change in alpha diversity can be found by differentiating 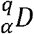 with respect to time. This leads to an expression that depends on changes in the relative abundance of each deme (Appendix S1), though it still loses information on changes in the total number of individuals across the metacommunity.

Dispersal can further obscure the effects of biotic interactions because changes in relative abundance in one community may be due to some combination of in situ growth and dispersal. Knowledge of dispersal can be used to tease apart these effects. Established tools can tease apart the contributions of in situ growth versus dispersal (Kerr and Godfrey-Smith 2009), while uncertainty about the extent of dispersal can be incorporated, say by simulating (Godsoe et al. 2021). We can learn about the changes in relative abundances of each deme due to in situ growth. As with our original model, analyses of changes in relative abundances still miss information on *N*.

Beta diversity is also insensitive to changes in *N*. This is because beta diversity is defined as the ratio of gamma to alpha diversity (Jost 2007), neither of which contain information on N (See Appendix S1). This conclusion also applies to some pairwise measurements of species beta diversity such as Jaccard, Sørensen, Horn and Morrista-Horn indices which are just monotonic transformations of Hill numbers (Jost 2007).

### Empirical analysis of diversity

Early experiments on *Drosophila* illustrate the gap between biotic interactions and changes in biodiversity. Ayala et al. (1973) presents data on competition between *D. pseudoobscura* and *D. willistoni* in cultures. In culture these two species coexist, but compete intensely (competition roughly halves the equilibrium density of both species). To measure the effect of competition Ayala et al. (1973) created replicated cultures of 19 different combinations of abundances of both species and then measured the short-term change in the number of individuals in each culture. We compare this analysis of competition to an analysis of diversity within each treatment group, and an analysis where total change across treatments is partitioned into contributions from alpha, beta and gamma diversity.

In many treatments, absolute abundances decline while relative abundances change weakly (Fig. 5 A). As a result, the change in diversity measurements within each locality Δ ^2^*D*, cannot be predicted using changes in total abundance (p=0.9968, *R*^2^=0, here ^2^*D* is local diversity and Δ denotes the difference between local diversity at the end versus start of the experiment). In one such treatment, the relative abundance of *D. willistoni* increased from 0.25 to 0.3 (blue arrow), while changes in absolute abundances are small. In a second example (red arrow) relative abundance remained 0.13 before and after treatment. Negligible change in relative abundance results in negligible change in diversity.

**Figure 4.**
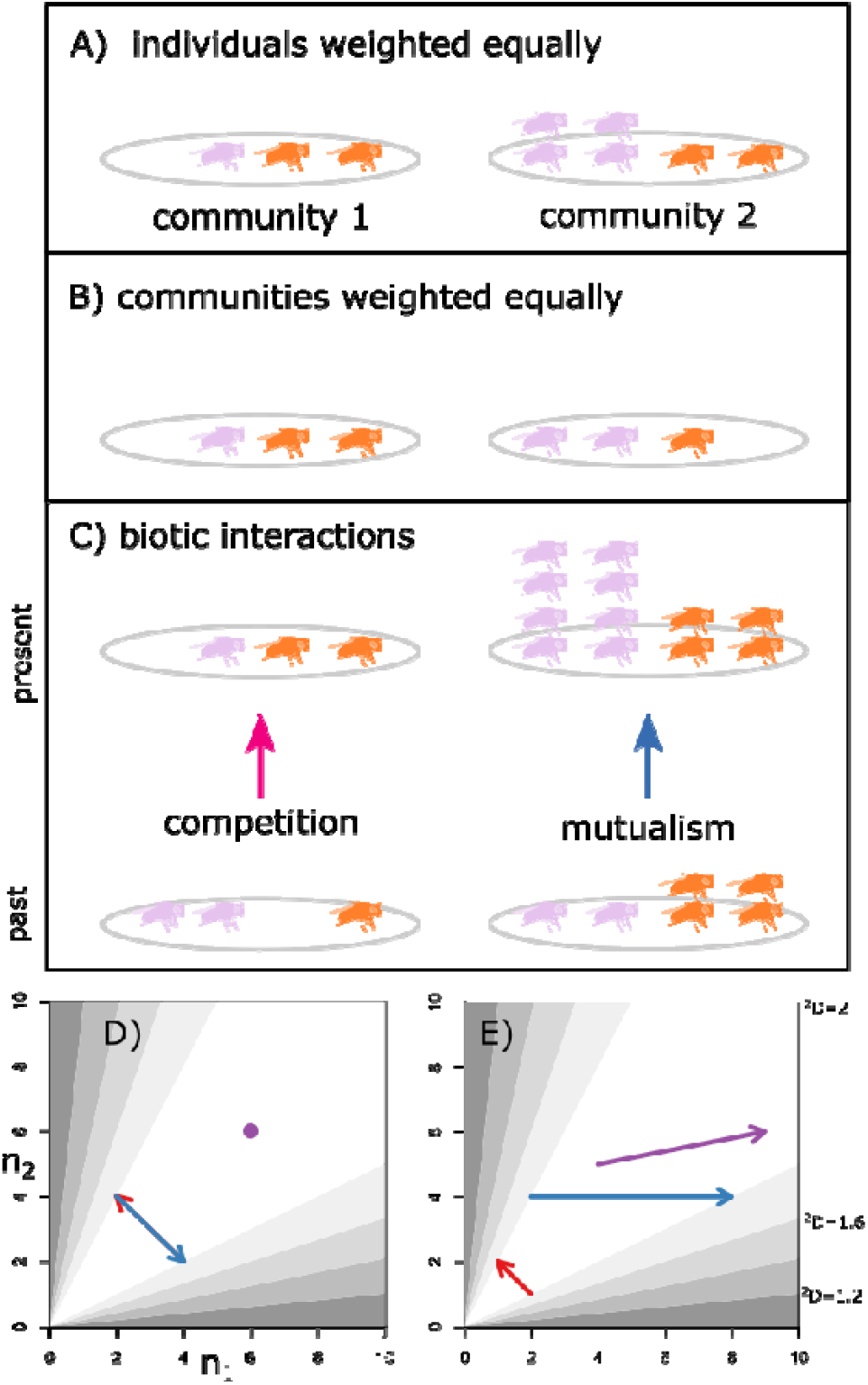
illustrates how diversity partitions can obscure effects of biotic interactions across a metacommunity. A) Illustrates a simplified metacommunity where community 1 has fewer individuals than community 2. B) For most Hill numbers the recommended procedure is to weight communities equally, we illustrate this here by setting the number of individuals to be the same in both communities (i.e. 3). In effect, this de-emphasizes the contribution of individuals in community 2. C) Illustrates the effects of biotic interactions across a metacommunity, with competition in community 1 reducing the relative abundance of purple flies, while mutualism in community 2 leads to increases in absolute abundances of both purple and orange flies and an increase in the relative abundances of purple flies. D) There is no change in diversity when communities are weighted equally. The changes within community 1 (red arrow) are counterbalanced by the changes in community 2 (blue arrow). Overall diversity remains unchanged (purple dot). D) illustrates changes in diversity when individuals are weighted equally; here gamma diversity changes (purple arrow) because the relative abundance of the purple flies across the metacommunity increases.

**Figure 5.**
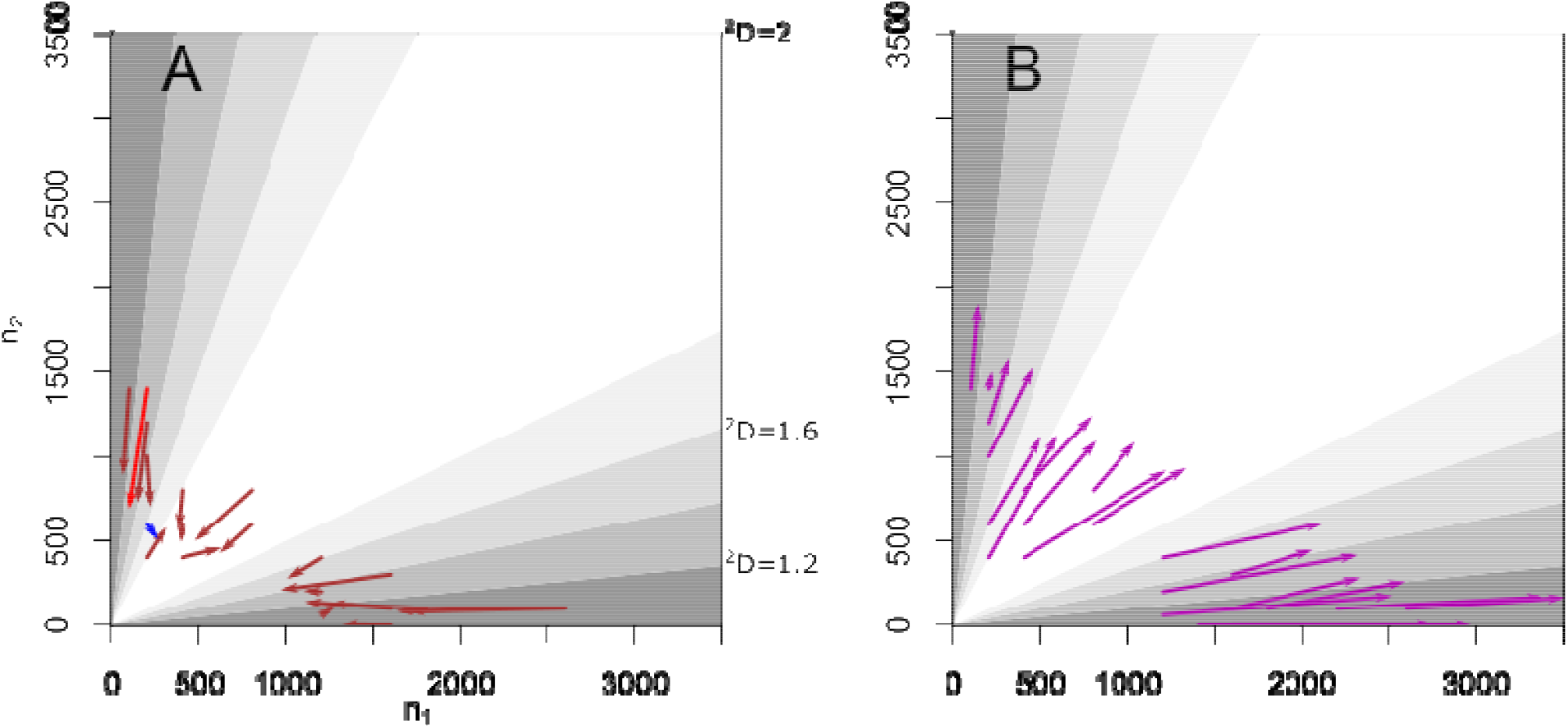
A) Population dynamics from competition experiments for two species of fruit flies *Drosophila willistoni* and *Drosophila pseudoobscura*. Each arrow represents one set of initial conditions with the tail showing the absolute abundances of species at the start of an experiment and the tip showing absolute abundances at the end of the experiment. The red arrow is represents experimental conditions where N declines but relative abundances and diversities remained unchanged. The blue arrow represents experimental conditions where change in N is far smaller but changes in relative abundance and diversity are larger. B) Illustrates an artificial dataset including simulated changes in absolute abundances across all communities. This artificial dataset leads to exactly the same changes in diversity as observed in the original dataset.

Across treatments the total number of flies declines by around 20 percent (Fig. 5 A) consistent with the high initial population densities and strong competition reported by Gilpin and Ayala (1973). Over the experiment, there is an increase in alpha diversity and a decrease in beta and gamma diversity, 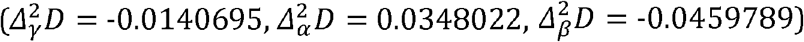. These results are consistent with the interpretation provided by (Gilpin and Ayala 1973), that the species compete intensely but different initial conditions converge on an equilibrium where the two species coexist. However, the observed changes in diversity are consistent with radically different ecological scenarios. We illustrate the insensitivity of diversity to total population with a thought experiment based on artificial dataset that exhibits population growth across all communities same diversity changes, but produces the same changes in diversity (Fig. 5 B). In this plot, the initial populations are unchanged but the total population size at the end of the experiment has been doubled across all treatments. This results in rapid population growth across the metacommunity. Though Fig. 5 B, relaxes the intense competition observed empirically, diversity changes are identical to those in (Fig. 5 A; 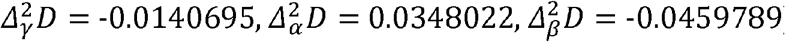).

## Discussion

We have proposed a synthesis of species interactions and diversity changes. This synthesis highlights how changes in diversity can capture some facets of species interactions, such as differences in per-capita growth rates among species. Other facets are lost such as shifts in *N*. For more than a century, ecologists have sought to relate biotic interactions and measurements of diversity (Barner et al. 2018; Michael 1920; Simha et al. 2022). Our work suggests that it is time to move past this debate and recognize that some questions about biotic interactions are unanswerable with data on biodiversity. We will highlight three potential mitigation strategies: (i) articulating the assumptions needed to link biotic interactions and diversity; (ii) deriving mechanistic models based on partitions of diversity change; (iii) using diversity measurement that maximize the information available. Ecologist’s ability to conserve biodiversity depends on reliable models (Socolar et al. 2016; Urban et al. 2016). We seek to improve these models by clarifying which information on biotic interactions is needed to improve our models.

A major insight from this synthesis is that changes in biodiversity are a poor surrogate for the effects of biotic interactions. This is because indices of diversity omit information on the total number of individuals in the metacommunity (*N*). When this information is missing, different biotic interactions are capable of producing comparable changes in diversity (Fig. 1, 3). To understand this mismatch, we tie together models of biotic interactions (Abrams 1987), relative abundance (Lion 2018), and diversity change (Frank and Godsoe 2020; Godsoe et al. 2021). However, our conclusions are limited to analyses of short-term dynamics. It is still possible that there exists some signature that can be detected with other sources of data. For example analyses of long-term dynamics over multiple generations (Schaffer 1981), or measurements of the similarity of species (Leinster and Cobbold 2012). However, we urge caution, biotic interactions lead to a diverse array of ecological dynamics and there can be considerable overlap in the dynamics produced by each type of interaction (Holland and DeAngelis 2009; Holland and DeAngelis 2010; Mohd et al. 2016). Moreover, additional sources of information like trait or phylogenetic similarity sometimes have unexpected effects on biotic interactions. For example, it is commonly assumed that increasing similarity among competitors strengthens the effects of competition (HilleRisLambers et al. 2012), but an analysis using annual plants in California found that similarity measured using single traits decreases the strength of competition (Kraft et al. 2015).

Over long timescales, biotic interactions may drive speciation (Nosil 2012; Yoder and Nuismer 2010). For example, mutualistic interactions among yuccas and yucca moths leads to co-speciation, with each partner matching the morphology of the other (Godsoe et al. 2008). A process which has taken hundreds of thousands if not millions of years (Smith et al. 2008; Smith et al. 2011). Effects of biotic interactions on speciation may be detectable using analyses of phylogenies (Nuismer and Harmon 2015; Pigot and Tobias 2013; Yoder and Nuismer 2010), but these longer-term trends are beyond the scope of the current manuscript.

Many previous dynamic models of biodiversity make it difficult to detect dependence on absolute abundances because they assume that community dynamics are zero-sum, implying the total number of individuals in a metacommunity is constant (Hubbell 2001). Analyses of species coexistence focusing on frequency dependence also make this zero sum assumption (Godwin et al. 2020). This zero sum assumption is only valid for some models, notably Lotka-Volterra competition among similar species (Lion 2018; Mallet 2012), it does not typically emerge from models of other interaction types (Holland and DeAngelis 2009; Holland and DeAngelis 2010).

There is compelling evidence that community assembly is non-zero sum in nature. Notably, diversity productivity experiments have been designed to tease apart selection effects (which are zero sum) from non-zero sum effects such as niche complementarity and facilitation (Cardinale et al. 2012; Loreau and Hector 2001). A sizeable proportion of these effects are due to the non-zero sum dynamics (Cardinale et al. 2012). Worse, systems might change from zero sum to non-zero sum as we move from one community to another. Interactions among plants in low elevation tend to be competitive (and may be close to zero sum) while interactions in high elevation sites are more likely to be facilitative (Callaway et al. 2002; Louthan et al. 2015). Switches between one interaction type and another are common in nature (Chamberlain et al. 2014).

One way to mitigate these problems is by modelling mechanisms that change relative abundances rather than absolute abundances. For example, Vellend (2010) argues that total change in diversity should be divided into four fundamental mechanisms selection, drift, immigration and speciation. In this framework, salient effects of biotic interactions are captured by selection and drift. Several groups have developed tools to quantify these mechanisms in long-term dataset using partitioning techniques generate models that start with the measurement of interest (i.e. diversity) then ask how a small amount of change in each type of organism will change diversity. For example, Tatsumi et al. (2021) showed that changes in Jaccard similarity, a measure of beta diversity can be partitioned into consequences of extinctions and colonization’s. When applied to long-term forest plots they were able to show that extinction events and colonization events roughly balanced out, leading negligible change in Jaccard similarity. Similarly (Godsoe et al. 2021) used a partitioning approach to show that changes in diversity are often due to the success of rare species relative to common ones. This approach outlines each of the mechanisms described in (Vellend 2010) can be analysed, including speciation, drift and immigration. Extensions to Hill numbers and beta diversity can be found in (Frank and Godsoe 2020; Godsoe et al. 2022). Though the development of partitions for diversity change are relatively recent, these techniques draw on a rich literature of partition methods developed for other problems in ecology and evolution (Collins and Gardner 2009; Fox 2016; Loreau and Hector 2001).

Given the difficulties we identify some might wonder if we need measures that can detect biotic interactions directly. The answer is this has been tried, repeatedly, over a more than a century. See Blanchette et al. (2020) for an up to date summary of dozens of such proposals ranging from (Forbes 1907) up to the present day. Blanchette et al. note that the accuracy of these methods tested repeatedly (Barner et al. 2018; Brazeau and Schamp 2019; Freilich et al. 2018; Thurman et al. 2019) with tests finding that “current methods are generally inaccurate, and thus, the spatial associations detected are poor proxies for biotic interactions”. In view of this, we believe that progress in the next century will not require more solutions. What is needed is a better understanding of the problem.

We have shown that there is a gap between traditional models of community assembly and change in modern measurements of biodiversity. This may be disconcerting to who might expect a synthesis of biotic interactions and biodiversity to solve problems and lead to new methods to detect biotic interactions. We disagree, in our view synthesis in biology typically clarify the connections and purposes of different approaches (Lion 2018). This is what equation 4 does, by explicitly connecting changes in diversity to changes in relative abundances generated by biotic interactions. Biodiversity misses changes in absolute abundances, which can be crucial to traditional models of community dynamics. Our work suggests that previous efforts to identify biotic interactions using short-term observations of biodiversity are misguided; since any observation will be compatible with multiple contradictory mechanisms. The gap we identify can be mitigated by explicitly modelling mechanisms that shape diversity (such as shifts in relative abundances) rather than modelling mechanism that have traditionally interested ecologists such as competition.

## Acknowledgements

We thank wint for pointing out “theres [sic] actually zero difference between good & bad things”. Helpful comments were provided by Jennifer Bufford, Roseanna Gamlen-Greene, Andrew Letten, Zach Marion and Dr. Tim Poisot, and one anonymous reviewer. We thank the Society of Population Ecology, whose 2019 conference stimulated the initial idea for this project. This paper was only possible because of virtual meetings between Aotearoa and Japan. Thus, in spite of our better judgement, we thank Google Jamboard, Zoom and Microsoft Teams.

---

**Table 1.**
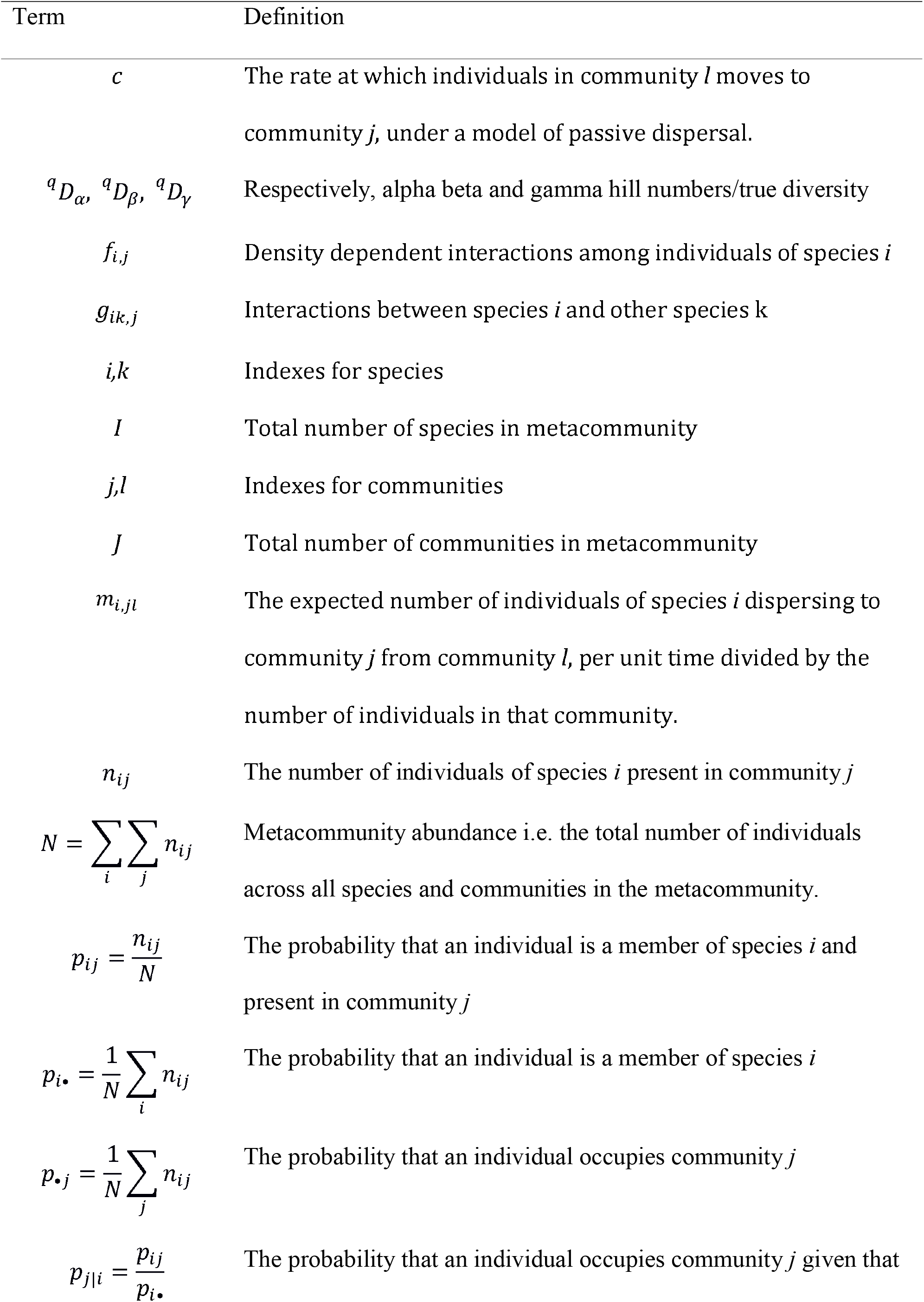
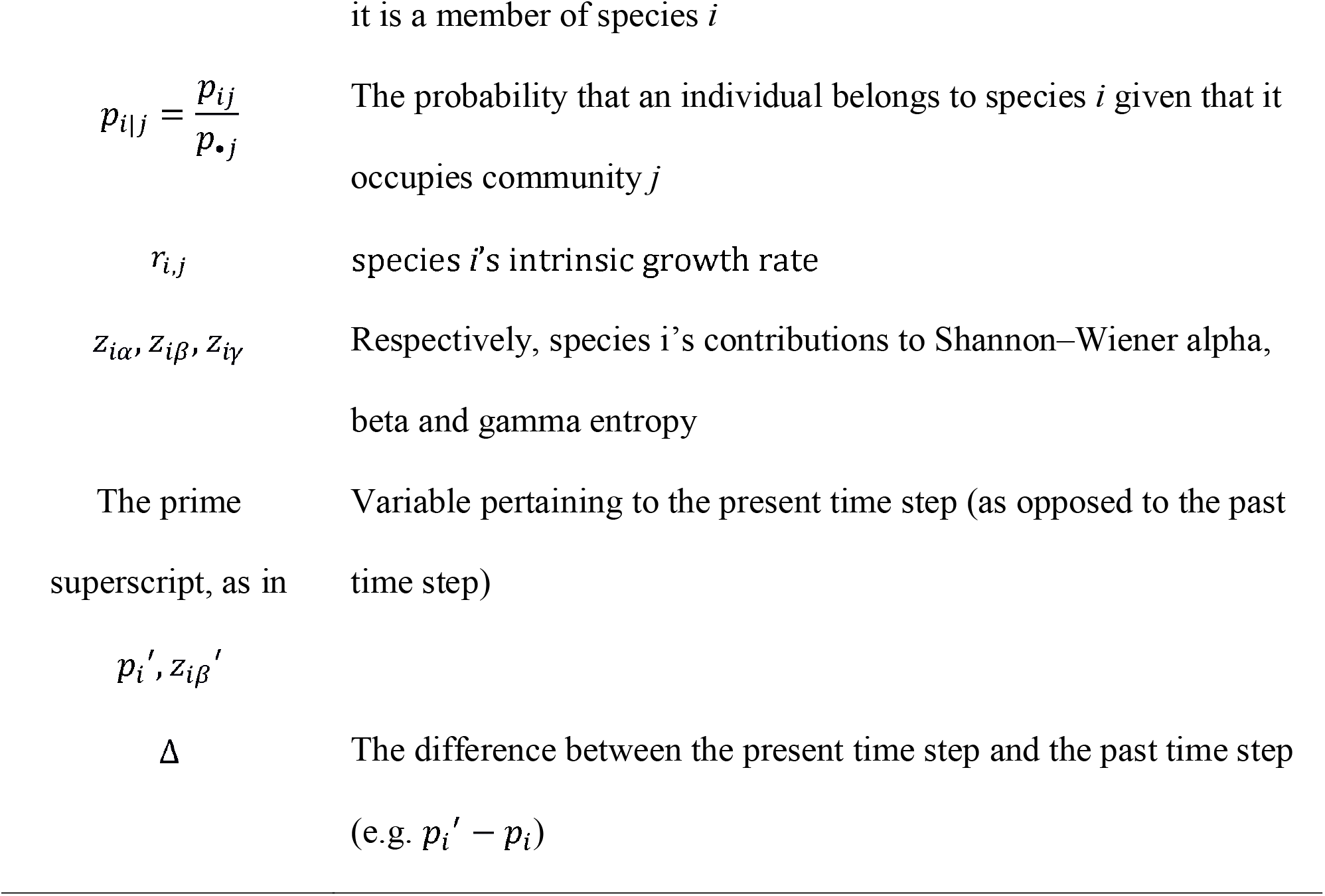
list of terms used in the main text.

~~~
---
title: “Appendix S2 for: Hill numbers unify interactions and diversity”
author: “William Godsoe”
date: “17/01/2022”
output:
html_document: default
 word_document: default
 pdf_document: default
---
This RMD file was used to generate drafts of the analyses presented in the draft manuscript as of January 2022. Code was last run in R 4.0.2.
note the code was written using the d function in the vegetarian package, to compute diversity statistics. As of August 2021 this package has been removed from CRAN. Archived versions of the function are available https://cran.r-project.org/web/packages/vegetarian/index.html
Contact William Godsoe for queries william.godsoe@lincoln.ac.nz to generate figure 5 you will need to download a dataset of fly abundances found in the file “AyalaTable1.csv”. Then modify the read.csv file commented out on line 41 to the new path. Finally change the header at L874 from eval=F to eval=T.
‘‘‘{r setup, include=FALSE}
knitr::opts_chunk$set(echo = TRUE)
require(phaseR)
require(deSolve)
require(RColorBrewer)
require(vegetarian)
require(ggplot2)
require(reshape2)
require(plyr)
require(dplyr)
require(tidyr)
require(vegan)
require(knitr)
Ayala.competition <- read.csv(“empirical phase
portraits/AyalaTable1.csv”)
#suggested colour palette
#colfunc <- colorRampPalette(brewer.pal(9,”Blues”)[c(8,2)])
colfunc <- colorRampPalette(grey(c(0.3,0.99)))
myScheme<-brewer.pal(8, “Accent”)
myGrey<-brewer.pal(8, “Greys”)[5:1]
myGrad<-brewer.pal(5,”GnBu”)
myGrad[1]<-”darkolivegreen” #helps richness show up better
LotVmod <- function (Time, State, Pars) {
 with(as.list(c(State, Pars)), {
  dx = r1*x-a11*x^2 +a12*y*x #note that interaction
  dy = r2*y+a21*x*y -a22*y^2
  return(list(c(dx, dy)))
 })
}
diCalcq<-function(p,qrare=1){
 divGrad1<-p
 for(i in 1:dim(p)[1]){
  for(j in 1:dim(p)[2]){
  divGrad1[i,j]<-d(c(p[i,j],1-p[i,j]), q=qrare) #Frequencies of two species
  }
 }
 return(divGrad1)
}
computeProbs<-function(x,y){
 p1<- x/(x+y)
}
calculateDiv<-function(comMat){
 tPoint<-comMat[1]
 comMat<-matrix(comMat[-1],2,2, byrow=T)
 #weights<-colSums(comMat)/sum(comMat)
 #note this has been corrected, i believe that rowsums is correct
 #for weighting communities by the # of individuals they contain weights<-rowSums(comMat)/sum(comMat)
 alpha=d(comMat,q=2,lev=“alpha”, wts=weights)
 beta=d(comMat,q=2,lev=“beta”, wts=weights)
 gamma=d(comMat,q=2,lev=“gamma”, wts=weights)
 return(c(time=tPoint,alpha=alpha,beta=beta,gamma=gamma))
}
plotODE<-function(Pars1, State,tmax=20,byLength=1, bylab=““,
Xlab=“time”,FigLab=““,figPos=1.9){
 Time <- seq(0, tmax, by = byLength)
 out <- as.data.frame(ode(func = LotVmod, y=State, parms = Pars1, times = Time))
 Dq2<-apply(out[,-1], 1, d, q=2)
 plot(Dq2∼Time, type=“l”, col=myGrad[3], lwd=2, xlab=Xlab,ylab=““)
 text(tmax*0.8, tail(Dq2)[1]-0.025, “q=2”, cex=1, col=“black”)
 Dq0<-apply(out[,-1], 1, d, q=0)
 points(Dq0∼Time, type=“l”, col=myGrad[1], lwd=2)
 text(tmax*0.8, tail(Dq0)[1]-0.025, “q=0”, cex=1, col=“black”)
 Dq1<-apply(out[,-1], 1, d, q=1)
 points(Dq1∼Time, type=“l”, col=myGrad[2], lwd=2)
 text(tmax*0.8, tail(Dq1)[1]-0.025, “q=1”, cex=1, col=“black”)
 Dq100<-apply(out[,-1], 1, d, q=100)
 points(Dq100∼Time, type=“l”, col=myGrad[4], lwd=2)
 text(tmax*0.8, tail(Dq100)[1]-0.025, “q=100”, cex=1, col=“black”)
 text(figPos,1.9, FigLab, cex=2, col=“black”)
 #text(tmax*0.8, 1.97, “q=0”, cex=1, col=“black”)
 #text(tmax*0.8, 1.78, “q=1”, cex=1, col=“black”)
 #text(tmax*0.8, 1.64, “q=2”, cex=1, col=“black”)
 #text(tmax*0.8, 1.38, “q=100”, cex=1, col=“black”)
}
hillFinder<-function(x){
 #this function takes the desired value for
 #hill# 2 for a community of two species
 (x-sqrt(-(x-2)*x))/(2*x)
}
drawHillGrad<-function(Xlab=““,xmax=1){
 plot(0:1,0:1,type=“n”,xlim = c(0, xmax),
  ylim = c(0, xmax),
  xaxs = “i”,
  yaxs = “i”,
  xlab=bquote(“n”[1]),
  ylab=bquote(“n”[2]))
 divPts<-seq(1.2,2, 0.2)
 slopeHolder<-NA
 for(HillV in 1:length(divPts)){
 relAbund<-hillFinder(divPts[HillV])
 slope<-(1-relAbund)/relAbund
 #abline(a=0,b=slope)
 #abline(a=0,b=1/slope)
 slopeHolder[HillV]<-slope
#original line was(I think)
#mtext(bquote(““[gamma]^”2”*”D” == .(divPts[HillV])),
mtext(bquote(““^”2”*”D” == .(divPts[HillV])),
  side=4,
  at=(1/slope)*xmax,
  cex=0.5,
  las=1)
}
extent<-xmax
polygon(x=c(0,
            0,
            extent),
        y=c(0,
            extent,
            slopeHolder[1]*extent),
        col=myGrey[1],
        lty=0)
polygon(x=c(0,
            extent,
            extent),
        y=c(0,
            0,
            1/slopeHolder[1]*extent),
        col=myGrey[1],
        lty=0)
for(i in 2:length(slopeHolder)){
 polygon(x=c(0,
            extent,
            extent),
        y=c(0,
            slopeHolder[i-1]*extent,
            slopeHolder[i]*extent),
        col=myGrey[i],lty=0)
 polygon(x=c(0,
            extent,
            extent),
         y=c(0,
            1/slopeHolder[i-1]*extent,
            1/slopeHolder[i]*extent),
         col=myGrey[i],lty=0)
}
}
### Added for figure 1
computeProbs<-function(x,y){
 p1<- x/(x+y)
}
computeN1<-function(x,y){
 p1<- x
}
computeN2<-function(x,y){
 p1<- y
}
drawArrows<-function(){
  arrows(Npts[1,]$N1,
         Npts[1,]$N2,
         Npts[2,]$N1,
         Npts[2,]$N2,
         length=0.1, col=annotCol[1], lwd=2)
  arrows(Npts[1,]$N1,
         Npts[1,]$N2,
         Npts[3,]$N1,
         Npts[3,]$N2,
         length=0.1, col=annotCol[1], lwd=2)
 arrows(Npts[1,]$N1,
         Npts[1,]$N2,
         Npts[4,]$N1,
         Npts[4,]$N2,
         length=0.1, col=annotCol[1], lwd=2)
 #i’m hiding this one for now
 arrows(Npts[1,]$N1,
         Npts[1,]$N2,
         Npts[5,]$N1,
         Npts[5,]$N2,
         length=0.1, col=NA,lwd=3)
 text(3,1, “1”)
 text(6,3.5, “2”)
 text(3,3.5, “3”)
}
drawRelAbund<-function(Xlab=““,xmax=1){
 extent<-xmax
 plot(0:1,0:1,type=“n”,xlim = c(0, xmax),
   ylim = c(0, xmax),
   xaxs = “i”,
   yaxs = “i”,
   xlab=Xlab)
 relGrad<-7
 relGrey<-brewer.pal(8, “Greys”)[1:relGrad]
 divPts<-seq(0,1,length.out=relGrad)
 divPts<-divPts/(1-divPts)
 for(i in 2:(relGrad-1)){
  polygon(x=c(0,
              extent,
              extent),
          y=c(0,
              divPts[i-1]*extent,
              divPts[i]*extent),
          col=relGrey[i],
          lty=0)
 }
 #add final polygon
 polygon(x=c(0,
             extent,
             0),
         y=c(0,
             divPts[i]*extent,
             extent),
         col=relGrey[relGrad],
         lty=0)
 abline(v=0)
}
labelK<-function(pars1){
 #mtext(bquote(“K”[1]),side=1, at=pars1[“r1”]/pars1[“a11”],cex=1,line=2)
 #mtext(bquote(“K”[2]), side=2, at=pars1[“r2”]/pars1[“a22”],
cex=1,line=2)
 abline(v=pars1[“r1”]/pars1[“a11”],col=brewer.pal(3, “Accent”)[2],lwd=2, lty=6)
 abline(h=pars1[“r2”]/pars1[“a22”],col=brewer.pal(3, “Accent”)[3],lwd=2, lty=6)
}
#this code is used to create a color gradient
#This is the code for the lotka-volterra model.
Pars <- c(r1=1, a11=1, a12=.9,
                r2=1, a21=.9, a22=1)
State <- c(x = 1, y = 1)
Time <- seq(0, 100, by = 1)
kcalcN<-2*Pars[‘r1’]/Pars[‘a11’]
yini=c(x=0.95,y=0.05)
annotCol<-brewer.pal(8,”Accent”)
‘‘‘
# Figure 1: Introducing diversity phase portraits
‘‘‘{r pretty1, cache=T, fig.width=6, fig.height=2, echo=F, cache=F}
N1 <- N2 <- seq(0, 10, length.out = 200)
p <- outer(N1, N2, computeProbs)
p[is.na(p)]<-0 #at 0,0 this returns NaN, to reduce problems downstream i set this to 0.
N1M <- outer(N1, N2, computeN1)
N2M <- outer(N1, N2, computeN2)
N <- N1M+N2M
par(mfrow=c(1,3), mar=c(4,3,1,2))
plot(1,1, type=“n”, ylim=c(0,10),xlim=c(0,10),xlab=“N1”, ylab=“N2”,
   xaxs = “i”,
   yaxs = “i”,
   main=“Absolute abundance”)
for(i in seq(0,10, by=2)){
 abline(h=i, col=“grey”)
 abline(v=i, col=“grey”)
}
Npts<-data.frame(N1=c(4,2,8,2,3), N2=c(2,1,4,4,3))
drawArrows()
points(Npts[1,], pch=16, cex=2)
points(Npts[2,], pch=16)
points(Npts[3,], pch=16)
points(Npts[4,], pch=16)
#points(Npts[5,], pch=16)
text(1, 9, “A”, cex=2, col=“black”)
#plot(1,1, type=“n”, ylim=c(0,10),xlim=c(0,10),xlab=“N1”, ylab=“N2”,
#      xaxs = “i”,
#     yaxs = “i”,
 #    main=“Relative abundance”)
drawRelAbund(xmax=10)
title(“Relative Abundance”)
drawArrows()
points(Npts[1,], pch=16, cex=2)
points(Npts[2,], pch=16)
points(Npts[3,], pch=16)
points(Npts[4,], pch=16)
#points(Npts[5,], pch=16)
text(1, 9, “B”, cex=2, col=“White”)
#plot(1,1, type=“n”, ylim=c(0,10),xlim=c(0,10),xlab=“N1”, ylab=“N2”,
#image(x=N1, y=N2, z = diCalcq(p,2),
#     useRaster = TRUE, add=T,
#      col=colfunc(10), xlab=“N1”, ylab=“N2”)
drawHillGrad(xmax=10)
drawArrows()
title(“Diversity”)
points(4,2, pch=16, cex=2)
points(2,1, pch=16)
points(8,4, pch=16)
points(2,4, pch=16)
#points(3,3, pch=16)
text(1, 9, “C”, cex=2, col=“white”)
#modifies the default image code in R to compute proportions.
‘‘‘
# Figure 2: see bottom of document
# Figure 3: simulations of biotic interactions
‘‘‘{r biSim, echo=F,cache=T,cache=F, fig.width=5, fig.height=9}
par(mfrow=c(4,2), mar=c(4,5,1,1))
ParsN <- c(r1=.25, a11=1, a12=0,
              r2=.49, a21=0, a22=1)
drawHillGrad()
labelK(ParsN)
title(“No interaction (neutralism)”)
bah<-trajectory(LotVmod,
           y0          = yini,
           tlim        = c(0, 30),
           parameters  = ParsN,
           system      = “two.dim”, lwd=3, col=myScheme[1])
points(yini[1],yini[2], cex=2, pch=16)
text(0.1, 0.9, “A”, cex=2)
plotODE(ParsN,yini, 25, 0.1, Xlab=“Time”,FigLab=“B”,figPos = 1.9)
ParsC <- c(r1=.5, a11=1, a12=-.9,
                 r2=.515, a21=-.9, a22=1)
drawHillGrad()
title(“Competition”)
labelK(ParsC)
bah<-trajectory(LotVmod,
           y0          = yini,
           tlim        = c(0, 300),
           parameters  = ParsC,
           system      = “two.dim”, lwd=3, col=myScheme[1])
points(yini[1],yini[2], cex=2, pch=16)
text(0.1, 0.9, “C”, cex=2, col=“black”)
plotODE(ParsC, yini, 400,FigLab=“D”,figPos = 1.9)
#############################Part 2 mutualism###############
ParsM <- c(r1=.35, a11=1, a12=.1,
                 r2=.51, a21=.8, a22=1)
drawHillGrad()
labelK(ParsM)
title(“Mutualism”)
bah<-trajectory(LotVmod,
            y0          = yini,
            tlim        = c(0, 30),
            parameters  = ParsM,
            system      = “two.dim”, lwd=3, col=myScheme[1])
points(yini[1],yini[2], cex=2, pch=16)
text(0.1, 0.9, “E”, cex=2, col=“black”)
plotODE(ParsM, yini,15,0.1, FigLab=“F”,figPos = 1.9)
##################part 3 predator prey#####################
ParsP <- c(r1=.5, a11=1, a12=-.4,
               r2=.44, a21=.4, a22=1)
drawHillGrad()
labelK(ParsP)
title(“Predation”)
bah<-trajectory(LotVmod,
           y0          = yini,
           tlim        = c(0, 30),
           parameters  = ParsP,
           system      = “two.dim”, lwd=3, col=myScheme[1])
points(yini[1],yini[2], cex=2, pch=16)
text(0.1, 0.9, “G”, cex=2)
plotODE(ParsP,yini, 25,0.1, FigLab=“H”,figPos = 1.9)
‘‘‘
# Figure S2: metacommunity simulations
‘‘‘{r metacomSim, echo=F,cache=F, fig.width=4, fig.height=8,cache=F}
comCol<-brewer.pal(6,”Set1”)
ParsC1 <- c(r1=.5, a11=1, a12=-.9,
                   r2=.5, a21=-.9, a22=1)
####################part 2 overlay diversity###################
par(mfrow=c(4,2), mar=c(4,3,1,1))
yini1<-c(x=0.05,y=0.01)
yini2<-c(x=0.01,y=0.05)
drawHillGrad(Xlab=“A”)
#Lvnull <- nullclines(LotVmod,
#                  xlim      = c(0.01, 1),
#                  ylim        = c(0.01, 1),
#                  parameters  = ParsC,
#                  points      = 500,
#                  add         = TRUE,
#                  lwd         =1,
#                  add.legend = FALSE,
#                  col=brewer.pal(3, “Accent”)[2:3],
#                  xlab=“N1”,
#                  ylab=“N2”)
bah1<-trajectory(LotVmod,
           y0          = yini1,
           tlim        = c(0, 15),
           parameters  = ParsC1,
           system      = “two.dim”, lwd=3, col=comCol[1])
bah2<-trajectory(LotVmod,
           y0          = yini2,
           tlim        = c(0, 15),
           parameters  = ParsC1,
           system      = “two.dim”, lwd=3, col=comCol[2])
text(0.2, 0.9, “A”, cex=2, col=“Black”)
points(yini1[1],yini1[2], cex=2, pch=16)
points(yini2[1],yini2[2], cex=2, pch=16)
bahSum<-cbind(time=bah1$t,
                    N1c1=bah1$x,
                    N2c1=bah1$y,
                    N1c2=bah2$x,
                    N2c2=bah2$y)
bahSumTot<-cbind(N1=bahSum[,2]+bahSum[,4],N2=bahSum[,3]+bahSum[,5])
points(N2∼N1,data=bahSumTot, col=comCol[4],lwd=3, type=“l”)
calculateDiv(bahSum[1,])
matrix(bahSum[1,-1],2,2)
divSum<-apply(bahSum[,],1,calculateDiv)%>%
 unlist()%>%
 t()%>%
 as.data.frame()
plot(divSum$alpha ∼ divSum$time.time, type=“l”, col=comCol[6], lwd=2, xlab=“Time”, ylim=c(0,2))
points(divSum$beta ∼ divSum$time.time, type=“l”, col=comCol[5], lwd=2)
points(divSum$gamma ∼ divSum$time.time, type=“l”, col=comCol[4], lwd=2)
text(1,head(divSum$alpha)[1]-0.1, bquote(““[alpha]^”2”*”D”),cex=1)
text(14,tail(divSum$beta)[1]-0.15, bquote(““[beta]^”2”*”D”),cex=1)
text(14,tail(divSum$gamma)[1]-0.15, bquote(““[gamma]^”2”*”D”),cex=1)
text(1, 0.25, “B”, cex=2)
#this code for multipart works, but it is stupid
#it only accepts integer counts, which doesn’t
#makes sense for simulated data
#multipart(matrix(bahSum[1,-1],2)*100, index=“renyi”, scales=2, nsimul=2)
yini3<-c(0.1,0.9)
drawHillGrad(Xlab=“C”)
#Lvnull <- nullclines(LotVmod,
#                     xlim         = c(0.01, 1),
#                      ylim          = c(0.01, 1),
#                      parameters    = ParsC,
#                      points        = 500,
#                      add           = TRUE,
#                      lwd           =1,
#                     add.legend    = FALSE,
#                     col=brewer.pal(3, “Accent”)[2:3],
#                      xlab=“N1”,
#                      ylab=“N2”)
bah1<-trajectory(LotVmod,
            y0         = yini3,
            tlim       = c(0, 15),
            parameters = ParsC1,
            system     = “two.dim”, lwd=3, col=comCol[1])
bah2<-trajectory(LotVmod,
            y0         = yini3[2:1],
            tlim       = c(0, 15),
            parameters = ParsC1,
            system     = “two.dim”, lwd=3, col=comCol[2])
text(0.2, 0.9, “C”, cex=2)
points(yini3[1],yini3[2], cex=2, pch=16)
points(yini3[2],yini3[1], cex=2, pch=16)
bahSum<-cbind(time=bah1$t,
                     N1c1=bah1$x,
                     N2c1=bah1$y,
                     N1c2=bah2$x,
                     N2c2=bah2$y)
bahSumTot<-cbind(N1=bahSum[,2]+bahSum[,4],N2=bahSum[,3]+bahSum[,5])
points(N2∼N1,data=bahSumTot, col=comCol[4],lwd=3, type=“l”)
calculateDiv(bahSum[1,])
divSum<-apply(bahSum,1,calculateDiv)%>%
 unlist()%>%
 t()%>%
 as.data.frame()
plot(divSum$alpha ∼ divSum$time.time, type=“l”, col=comCol[6], lwd=2,
xlab=“Time”, ylim=c(0,2))
points(divSum$beta ∼ divSum$time.time, type=“l”, col=comCol[5], lwd=2)
points(divSum$gamma ∼ divSum$time.time, type=“l”, col=comCol[4], lwd=2)
text(1,head(divSum$alpha)[1]-0.15, bquote(““[alpha]^”2”*”D”),cex=1)
text(1,head(divSum$beta)[1]+0.15, bquote(““[beta]^”2”*”D”),cex=1)
text(14,tail(divSum$gamma)[1]-0.15, bquote(““[gamma]^”2”*”D”),cex=1)
text(1, 0.25, “D”, cex=2)
#this code for multipart works, but it is stupid
#it only accepts integer counts, which doesn’t
#makes sense for simulated data
#multipart(matrix(bahSum[1,-1],2)*100, index=“renyi”, scales=2, nsimul=2)
#################eg 3##############33
drawHillGrad(xmax=2)
bah1<-trajectory(LotVmod,
            y0         = yini,
            tlim       = c(0, 25),
            parameters = ParsM,
            system     = “two.dim”, lwd=3, col=comCol[1])
bah2<-trajectory(LotVmod,
           y0          = yini,
           tlim        = c(0, 25),
           parameters  = ParsP,
           system      = “two.dim”, lwd=3, col=comCol[2])
text(0.2, 1.9, “E”, cex=2)
points(yini[1],yini[2], cex=2, pch=16)
bahSum1<-cbind(time=bah1$t,
                    N1c1=bah1$x,
                    N2c1=bah1$y,
                    N1c2=bah2$x,
                    N2c2=bah2$y)
bahSumTot1<-cbind(N1=bahSum1[,2]+bahSum1[,4],N2=bahSum1[,3]+bahSum1[,5])
points(N2∼N1,data=bahSumTot1, col=comCol[4],lwd=3, type=“l”)
divSum<-apply(bahSum1,1,calculateDiv)%>%
 unlist()%>%
 t()%>%
 as.data.frame()
plot(divSum$alpha ∼ divSum$time.time,
      type=“l”,
      col=comCol[6],
      lwd=2,
      xlab=“Time”,
     # tail(divSum$alpha)[1],
      ylim=c(0,2))
points(divSum$beta ∼ divSum$time.time,
       type=“l”,
       col=comCol[5],
       lwd=2)
points(divSum$gamma ∼ divSum$time.time,
       type=“l”,
       col=comCol[4],
       lwd=2)
text(4,tail(divSum$alpha)[1]-0.15, bquote(““[alpha]^”2”*”D”),cex=1)
text(22,tail(divSum$beta)[1]-0.15, bquote(““[beta]^”2”*”D”),cex=1)
text(22,tail(divSum$gamma)[1]-0.15, bquote(““[gamma]^”2”*”D”),cex=1)
text(1, 0.25, “F”, cex=2)
#this code for multipart works, but it is stupid
#it only accepts integer counts, which doesn’t
#makes sense for simulated data
#multipart(matrix(bahSum[1,-1],2)*100, index=“renyi”, scales=2, nsimul=2)
‘‘‘
#### Figure 4 (in part): schematic of metacommunity change
‘‘‘{r metacommunityFig, fig.width=3, fig.height=10, echo=F}
par(mfrow=c(3, 1))
#plot(1,1, type=“n”, ylim=c(0,10),xlim=c(0,10),xlab=“N1”, ylab=“N2”,
#main=“Metacommunity”)
#image(x=N1, y=N2, z = diCalcq(p,2),
#    useRaster = TRUE, add=T,
#     col=colfunc(10), xlab=“N1”, ylab=“N2”)
drawHillGrad(xmax=10)
  arrows(Npts[1,]$N1,
         Npts[1,]$N2,
         Npts[2,]$N1,
         Npts[2,]$N2,
         length=0.1, col=comCol[1], lwd=3)
  arrows(Npts[1,]$N2,
         Npts[1,]$N1,
         Npts[2,]$N2,
         Npts[2,]$N1,
         length=0.1, col=comCol[2], lwd=3)
  #try to make combined arrow
 arrows(Npts[1,]$N1+Npts[1,]$N2,
         Npts[1,]$N2+Npts[1,]$N1,
         Npts[2,]$N1+Npts[2,]$N2,
         Npts[2,]$N2+Npts[2,]$N1,
         length=0.1, col=comCol[4], lwd=3)
  text(1, 9, “D”, cex=2, col=“white”)
  ###########################################################
 # plot(1,1, type=“n”, ylim=c(0,10),xlim=c(0,10),xlab=“N1”, ylab=“N2”,
#main=“Metacommunity”)
#image(x=N1, y=N2, z = diCalcq(p,2),
#      useRaster = TRUE, add=T,
#       col=colfunc(10), xlab=“N1”, ylab=“N2”)
    drawHillGrad(xmax=10)
#Npts<-data.frame(N1=c(4,2,8,2,3), N2=c(2,1,4,4,3))
  arrows(Npts[2,]$N1,
         Npts[2,]$N2,
         1,
         2,
         length=0.1, col=comCol[1], lwd=3)
  arrows(2,
         4,
         Npts[3,]$N1,
         Npts[3,]$N2,
         length=0.1, col=comCol[2], lwd=3)
      text(1, 9, “F”, cex=2, col=“white”)
  ###attempt at metacommunity arrow
   arrows(Npts[2,]$N1+2,
         Npts[2,]$N2+4,
         1+Npts[3,]$N1,
         2+Npts[3,]$N2,
         length=0.1, col=comCol[4], lwd=3)
   #####################
    # plot(1,1, type=“n”, ylim=c(0,10),xlim=c(0,10),xlab=“N1”, ylab=“N2”,
#main=“Metacommunity”)
#image(x=N1, y=N2, z = diCalcq(p,2),
#      useRaster = TRUE, add=T,
#       col=colfunc(10), xlab=“N1”, ylab=“N2”)
drawHillGrad(xmax=10)
#Npts<-data.frame(N1=c(4,2,8,2,3), N2=c(2,1,4,4,3))
 arrows(4,
        2,
        2,
        4,
        length=0.1, col=comCol[1], lwd=3)
 arrows(2,
        4,
        4,
        2,
        length=0.1, col=comCol[2], lwd=3)
    text(1, 9, “F”, cex=2, col=“white”)
 points(6, 6, col=comCol[4], cex=2, pch=16)
‘‘‘
# Figure 5: Empirical example
‘‘‘{r alayaNP, eval=T, fig.width=9, fig.height=5, echo=F}
N1<-Ayala.competition$willistoniE
N2<-Ayala.competition$pseudoobscuraE
changeN<-function(N1,N2,NF=2){
 N<-N1+N2
 P<-N1/(N1+N2)
 NM<-NF*N
 N1M<-NM*P
 N2M<-NM*(1-P)
 NnewN<-data.frame(N1M=N1M,N2M=N2M)
 return(NnewN)
 }
newPoints<-changeN(Ayala.competition$willistoniE,
          Ayala.competition$pseudoobscuraE, 2)
#recolored so that extreme cases are highlighted
Ayala.competition$deltaDCol<-”brown” colVec[1]
Ayala.competition$deltaDCol[3]<-”blue”
Ayala.competition$deltaDCol[6]<-”red”
par(mfrow=c(1,2))
#fruitPlot(qval=2, c(myScheme[7], myScheme[4], myScheme[5]),Nmax=3500)
drawHillGrad(xmax=3500)
text(400, 3300, “A”, cex=2)
segments(Ayala.competition$willistoniS,
       Ayala.competition$pseudoobscuraS,
       Ayala.competition$willistoniE,
       Ayala.competition$pseudoobscuraE,
        lwd=2,col=Ayala.competition$deltaDCol)
#points(Ayala.competition$willistoniE,
#       Ayala.competition$pseudoobscuraE, col=“red”,pch=16)
#fruitPlot(qval=2, c(myScheme[7], myScheme[4], myScheme[5]), Nmax=3500)
drawHillGrad(xmax=3500)
text(400, 3300, “B”, cex=2)
segments(Ayala.competition$willistoniS,
         Ayala.competition$pseudoobscuraS,
         newPoints[,1],
         newPoints[,2],
          lwd=2,col=“blue”) #col=Ayala.competition$deltaDCol)
‘‘‘
# Online table S1 parameters used in single species communities
‘r kable(rbind(ParsC, ParsM,ParsP, ParsN))
# Further details on simulations, nicely formatted
# Online table s2 parameters used in metacommunity
‘r kable(rbind(ParsC1, ParsC1, ParsM, ParsP))‘
# online table s2/s3 initial conditions used in metacommunity
‘r kable(rbind(yini1, yini2, yini3, yini3[2:1], yini, yini))‘
# Figure S1: plots of different Hill numbers
‘‘‘{r, echo=F, fig.width=7, fig.height=7, cache=T, eval=T}
#this function is broken, i’m not sure why and will need to fix when fresh
#takes a matrix of frequencies p and a diversity exponent q and returns diversity i.e. d()
par(mfrow=c(2,2), mar=c(4,4,3,3))
plotAltGrad<-function(Q, HillNum){
 plot(0:1,0:1,type=“n”,xlim = c(0, 1),
    ylim = c(0, 1),
    xaxs = “i”,
    yaxs = “i”,
    xlab=bquote(“n”[1]),
    ylab=bquote(“n”[2]),main=HillNum)
 image(z = diCalcq(p,Q),
     useRaster = T,
      col=colfunc(5), add=T)
}
plotAltGrad(Q=0, “Richness (q=0)”)
 abline(h=0,col=colfunc(5), lwd=5)
 abline(v=0,col=colfunc(5), lwd=5)
 text(0.1,0.9,”A”, cex=2)
plotAltGrad(Q=1, “Shannon (q=1)”)
 text(0.1,0.9,”B”, cex=2)
plotAltGrad(Q=2, “Simpsons (q=2)”)
text(0.1,0.9,”C”, cex=2)
plotAltGrad(Q=100, “Dominance (q=100)”)
 text(0.1,0.9,”D”, cex=2)
‘‘‘
‘‘‘{r convertdpdD, eval=F, fig.height=4, fig.width=5, echo=F}
arrowDraw<-function(Q,p,location,L,divOr){
  slope<- diff(Q)[location]/diff(p)[location]
  segments(p[location],Q[location],
       p[location]+L,Q[location]+slope*L, lwd=2)
# text(p[location], Q[location],
#     bquote(partialdiff^.(divOr)*”D”/partialdiff*”p”==.(slopeR)),
#     cex=1, col=“black”)
}
p<-seq(0,1,0.01)
p[2]<-0.0001
p[(length(p)-1)]<-0.9999
div<-cbind(p,(1-p))
q0<-apply(div,1,d,q=0)
q1<-apply(div,1,d,q=1)
q2<-apply(div,1,d,q=2)
q100<-apply(div,1,d,q=100)
plot(q0∼ p, type=“l”, col=myGrad[1], ylab=“diversity”, lwd=3,
ylim=c(1,2.2),
    xlab=expression(“p”[i]))
abline(h=1)
points(q1∼ p, type=“l”, col=myGrad[2], lwd=3)
#points(q2∼ p, type=“l”, col=myGrad[3], lwd=2)
points(q100∼ p, type=“l”, col=myGrad[4], lwd=3)
As<-60
AL<-0.05
#arrowDraw(q0,p,10,0.075)
arrowDraw(q1,p,As,AL,divOr=“1”)
#arrowDraw(q2,p,As,AL)
arrowDraw(q100,p,As,AL,divOr=“100”)
segments(p[As],1,
       p[As]+AL,1, lwd=2)
segments(p[As],2,
       p[As]+AL,2, lwd=2)
text(p[As], 1.1,
   expression(frac(“dp”[i],”dt”)==-0.05),
   cex=.75, col=“black”)
text(p[As], q0[As]+0.1,
   expression(frac(partialdiff^0*”D”,partialdiff*”p”[i])==0),
   cex=.75, col=“black”)
text(p[As]-0.05, q1[As],
   expression(frac(partialdiff^1*”D”,partialdiff*”p”[i])%∼∼%1),
   cex=.75, col=“black”)
text(p[As]-0.05, q100[As],
   expression(frac(partialdiff^100*”D”,partialdiff*”p”[i])%∼∼%3),
   cex=.75, col=“black”)
text(p[10], q0[10], expression(““^0*”D”))
text(p[10], q1[10], expression(““^1*”D”))
text(p[10], q100[10], expression(““^100*”D”))
p[As]
‘‘‘
# Figure 2: how changes in relative abundance shift into changes in diversity
‘‘‘{r convertdpdD1, fig.height=8, fig.width=8, echo=F}
arrowDraw<-function(Q,p,location,L,divOr){
   slope<- diff(Q)[location]/diff(p)[location]
   arrows(p[location],Q[location],
        p[location]+L,Q[location]+slope*L, lwd=3)
# text(p[location], Q[location],
#      bquote(partialdiff^.(divOr)*”D”/partialdiff*”p”==.(slopeR)),
#      cex=1, col=“black”)
}
p<-seq(0,1,0.01)
p[2]<-0.0001
p[(length(p)-1)]<-0.9999
div<-cbind(p,(1-p))
q0<-apply(div,1,d,q=0)
q1<-apply(div,1,d,q=1)
q2<-apply(div,1,d,q=2)
q100<-apply(div,1,d,q=100)
As<-41
AL<-0.1
par(mfrow=c(2,2), mar=c(4,4,2,0))
############# richness####################
plot(q0∼ p, type=“l”, col=“darkolivegreen”, ylab=“diversity”, lwd=3,
ylim=c(1,2.2),
            main=“Richness”,
      xlab=expression(“p”[i.]))
arrows(p[As],1,
      p[As]+AL,1, lwd=3)
#arrows(p[As],2,
#       p[As]+AL,2, lwd=4)
text(p[As], 1.1,
     expression(frac(“dp”[i.],”dt”)==0.05),
     cex=1, col=“black”)
text(p[As]-0.1, q0[As]+0.15,
 expression(frac(partialdiff[gamma]^0*”D”,partialdiff*”p”[i])==0),
 cex=1, col=“black”)
text(0.1, 2.1, “A”, cex=2)
##############Shannon ##################
plot(q1∼ p, type=“l”, col=myGrad[2], lwd=3,
     ylab=““,
     ylim=c(1,2.2),
     main=“Shannon”,
     xlab=expression(“p”[i]))
arrows(p[As],1,
       p[As]+AL,1, lwd=3)
arrowDraw(q1,p,As,AL,divOr=“1”)
text(p[As], 1.1,
     expression(frac(“dp”[i.],”dt”)==0.05),
     cex=1, col=“black”)
text(p[As]-0.1, q1[As],
   expression(frac(partialdiff[gamma]^1*”D”,partialdiff*”p”[i.])%∼∼%0.8),
   cex=1, col=“black”)
text(0.1, 2.1, “B”, cex=2)
##############Simpsons ##################
plot(q2∼ p, type=“l”, col=myGrad[3], ylab=“diversity”, lwd=3,
ylim=c(1,2.2),
     main=“Simpsons”,
     xlab=expression(“p”[i.]))
arrows(p[As],1,
       p[As]+AL,1, lwd=3)
arrowDraw(q2,p,As,AL,divOr=“2”)
text(p[As], 1.1,
     expression(frac(“dp”[i.],”dt”)==0.05),
     cex=1, col=“black”)
text(p[As]-0.1, q2[As],
     expression(frac(partialdiff[gamma]^2*”D”,partialdiff*”p”[i.])%∼∼%1.4),
    cex=1, col=“black”)
text(0.1, 2.1, “C”, cex=2)
###############Dominance#############
plot(q100∼ p, type=“l”, col=myGrad[4], lwd=3, ylim=c(1,2.2),
     main=“Dominance”,
     ylab=““,
     xlab=expression(“p”[i.]))
arrows(p[As],1,
       p[As]+AL,1, lwd=3)
arrowDraw(q100,p,As,AL,divOr=“100”)
text(p[As], 1.1,
     expression(frac(“dp”[i.],”dt”)==0.05),
     cex=1, col=“black”)
text(p[As]-0.15, q100[As],
expression(frac(partialdiff[gamma]^infinity*”D”,partialdiff*”p”[i.])%∼∼%2 .9),
    cex=1, col=“black”)
text(0.1, 2.1, “D”, cex=2)
‘‘‘
~~~

## Appendix S1 supplemental derivations

### Plots of absolute abundances versus diversity

In the main text we show the link between absolute abundances of two species and the Hill number associated with Simpson’s diversity (*q* = 2). In Fig. S1 we illustrate how individual Hill numbers are all diagonally symmetrical on this plot. The implication being that intuitions gained for Simpson’s diversity in the main text can easily be applied to other diversity indices.

**Figure S1.**
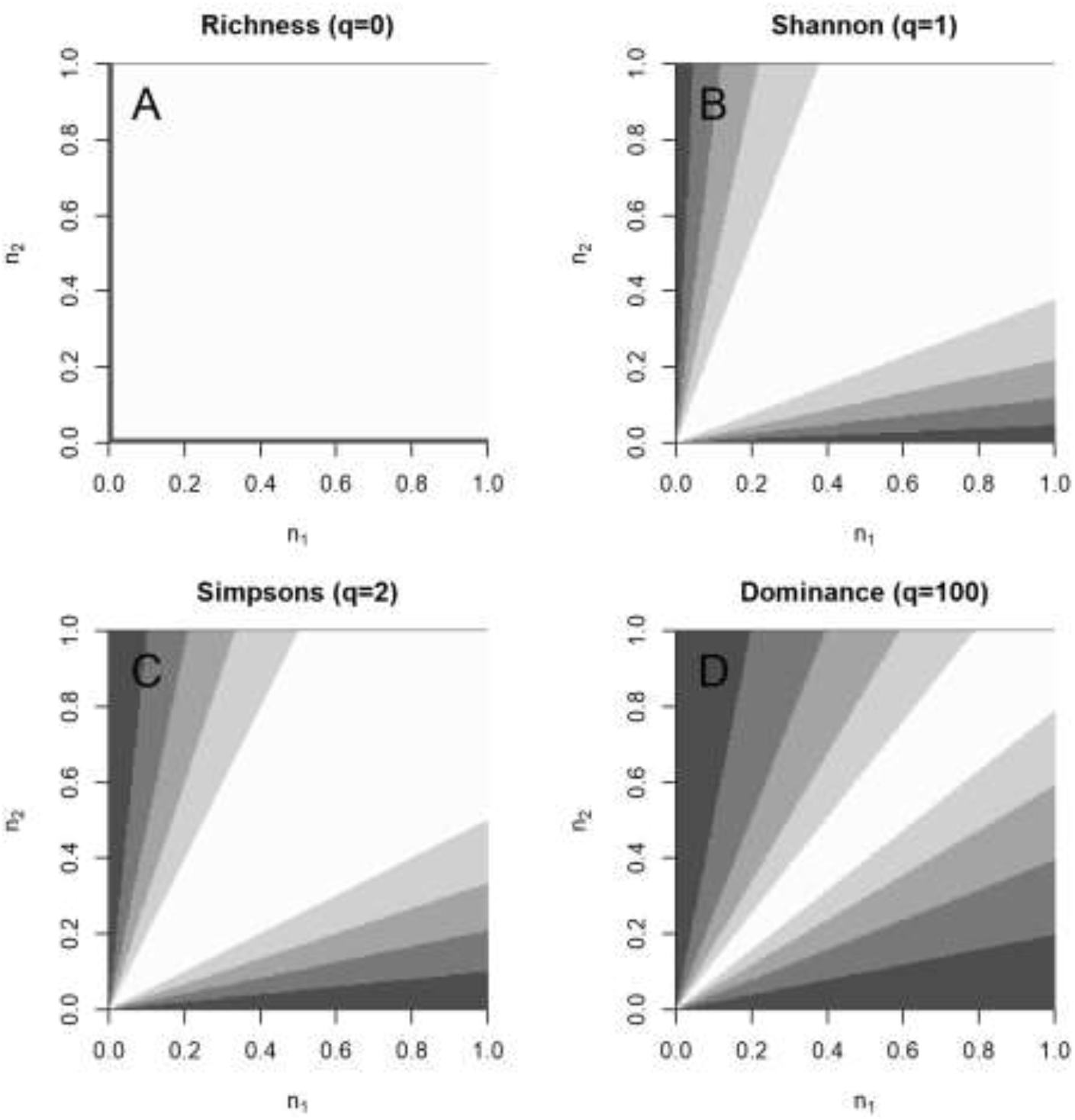
Overlays different diversities onto a plot of the absolute abundance of species 1 (*N*_1_) versus species 2 (*N*_2_). Plots are presented for different diversity orders from A) Richness (*q* = 0), B) Shannon Wiener (*q* = 1), C) Simpson’s (*q* = 2) and D) Dominance (large *q*; *q* = 100 in our case). All plots are diagonally symmetrical and range from a minimum of 1 to a maximum of 2. Note that richness only has a value of 1 when on the edges of the plot.

### Explanation of the per-capita effect of passive dispersal

In the main text we state, passive dispersal from community *l* into community *j* is modelled as 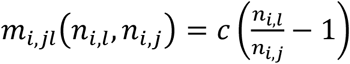. It is more intuitive and more common to present passive dispersal in a model of growth rate growth rate (Abrams and Wilson 2004). In our notation passive dispersal occurs when individuals of species *i* in community *l* emigrate to patch *j* at a rate *c* and individuals of species *i* in community *j* immigrate to patch l at a rate *c*. This leads to a model of change in *n*_*i,j*_ like:

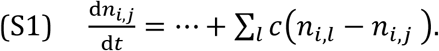

For emphasis, processes other than dispersal are here replaced by an ellipsis (…). Equation 1 in the main text is a model of per-capita change in populations. To convert equation S1 model to a per-capita rate of dispersal divide both sides by *n*_*i,j*_, giving:

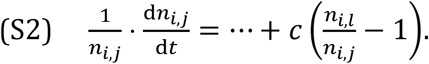

The form we use for passive dispersal in the main text.

### Expressions for alpha and beta diversity

Alpha diversity captures information on the relative abundance of each species within each community. Translated into the notation we use in the rest of the paper Jost (2007)’s expression for alpha diversity becomes:

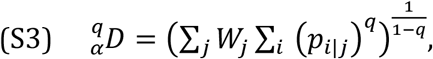

where *p*_*i*|*j*_ represents the relative abundance of species *i* in community 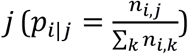. Each community is assigned a weight *W*_*j*_. Jost (2007) proposes a general expression for these weights 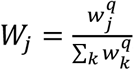, but recommends that all communities be weighted equally, for most diversity measurements. This is done by setting *W*_*j*_ = 1/*J*. The resulting expression for alpha diversity is:

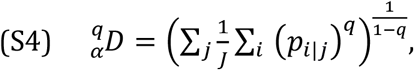

Differentiating this with respect to time leads to the following expression:

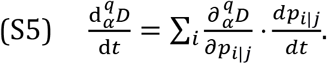

In words this means that change in alpha diversity can be found by computing the change in species’ frequencies within each community (*p*_*i*|*j*_) with respect to time 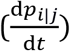, then translating changes in species’ frequencies into changes in diversity 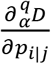 (*∂* denotes partial derivatives). Note that this expression implies that weights for each community stays constant over time. This requirement is consistent with the recommendation in Jost (2007) for most diversity indices, but as we have shown in the main text this choice can obscure important changes across the metacommunity (main text Fig. 4).

Since constraining the weights to be constant can lead to biases, we present an alternative expression for when the weightings vary over time. When this is the case, the total derivative for alpha diversity is given by:

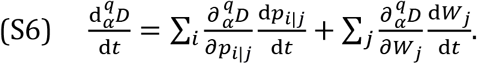

In this equation, the first term describes how the change in the frequency of species *i* in community *j*, changes alpha diversity over time. This is found by describing how alpha diversity changes over time 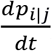, then converting these changes in *p*_*i*|*j*_ into changes in alpha diversity 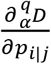. Alpha diversity reaches a maximum when all species in community *j* are equally abundant i.e. when 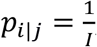, where *I* is the total number of species. Diversity increases monotonically as we move species *i*’s frequency towards this point (Patil and Taillie 1982).

The second term describes how alpha diversity changes as weights change. For example, it is possible to weight communities by the number of individuals they contain at a given time. Over time, the number of individuals can vary and hence the weights assigned to a given community can vary. Alpha diversity will increase when the weights assigned to communities with high local diversity increase relative to the weights assigned to communities with low local diversity.

The recommended approach to weight individuals equally for measurements of alpha diversity is to focus on the special case of Shannon entropy, found in the limit that *q* approaches 1:

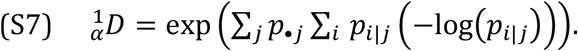

The expression for total derivative above is valid for Shannon diversity. To weight individuals equally, each community should be weighted by the probability that an individual in the metacommunity is found in that community: *W*_*j*_ = *p*_•*j*_. This leads to the following expression for diversity change:

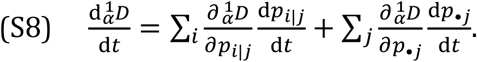

Note that changes in *p*_*i*|*j*_ and *p*_•*j*_ can be expressed in terms of changes in *p*_*ij*_. Beta diversity is the ratio of gamma diversity to alpha diversity, i.e.

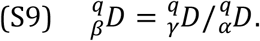

Therefore, the rate of change of beta diversity can be found using the quotient rule:

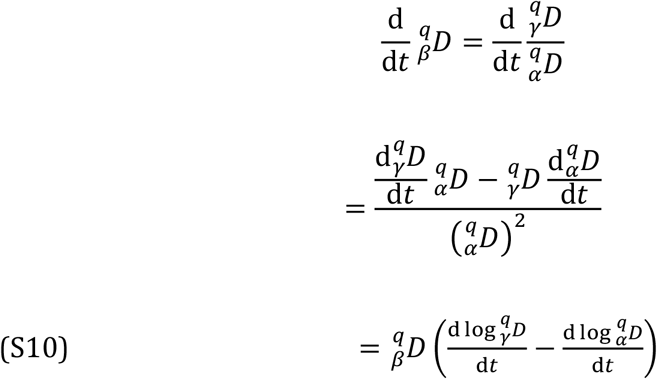

In words, the rate of change in beta diversity depends on the difference between change in log gamma diversity and change in log alpha diversity. No other sources of change are measured.

### Parameter values for simulations

**Table S1:**
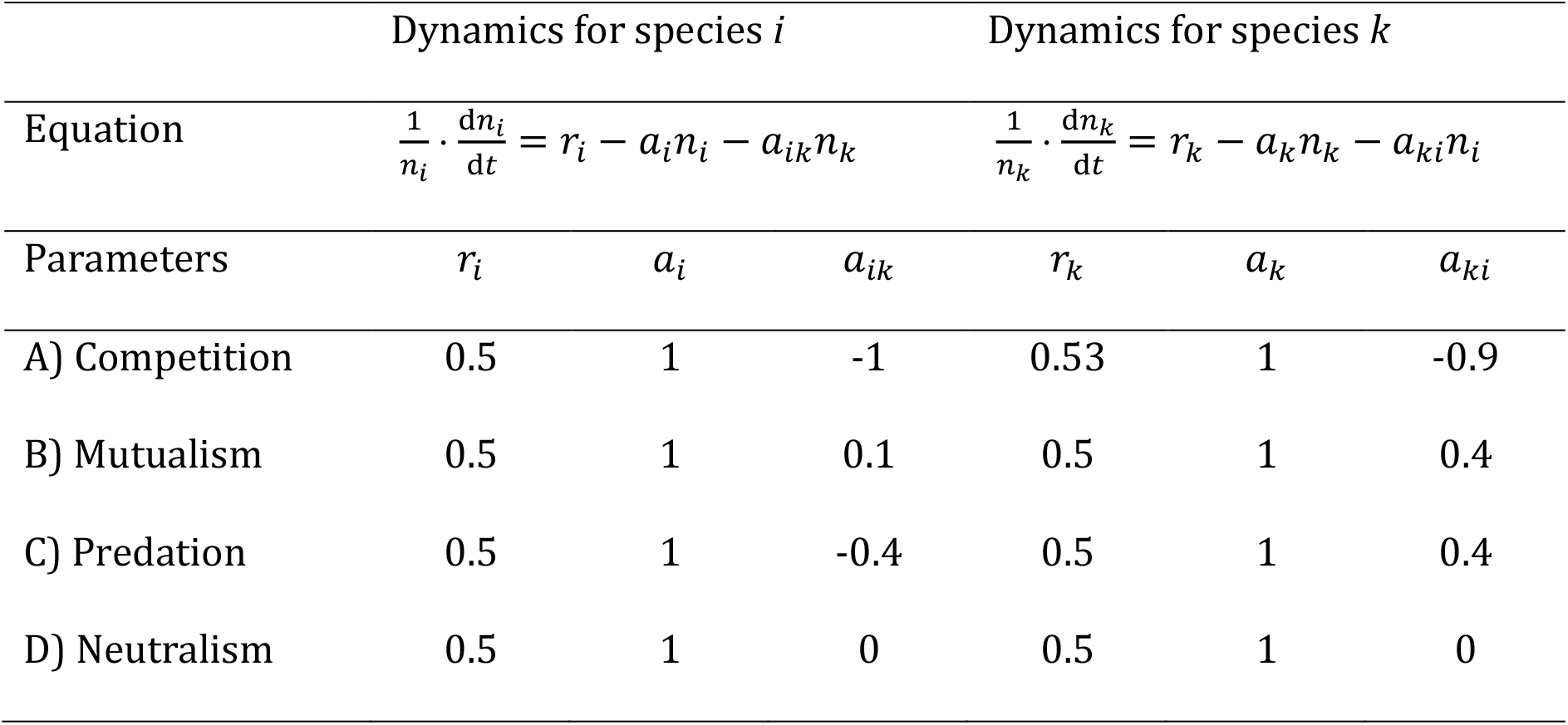
Parameter values used to simulate species interactions between species *i* and species *k* using a Lotka-Volterra model for simulations of Fig. 3 in the main text.

| | Dynamics for species $i$ | | | Dynamics for species $k$ | | |
| --- | --- | --- | --- | --- | --- | --- |
| Equation | $\frac{1}{n_i} \cdot \frac{dn_i}{dt} = r_i - a_i n_i - a_{ik} n_k$ | | | $\frac{1}{n_k} \cdot \frac{dn_k}{dt} = r_k - a_k n_k - a_{ki} n_i$ | | |
| Parameters | $r_i$ | $a_i$ | $a_{ik}$ | $r_k$ | $a_k$ | $a_{ki}$ |
| A) Competition | 0.5 | 1 | -1 | 0.53 | 1 | -0.9 |
| B) Mutualism | 0.5 | 1 | 0.1 | 0.5 | 1 | 0.4 |
| C) Predation | 0.5 | 1 | -0.4 | 0.5 | 1 | 0.4 |
| D) Neutralism | 0.5 | 1 | 0 | 0.5 | 1 | 0 |

**Figure S2:**
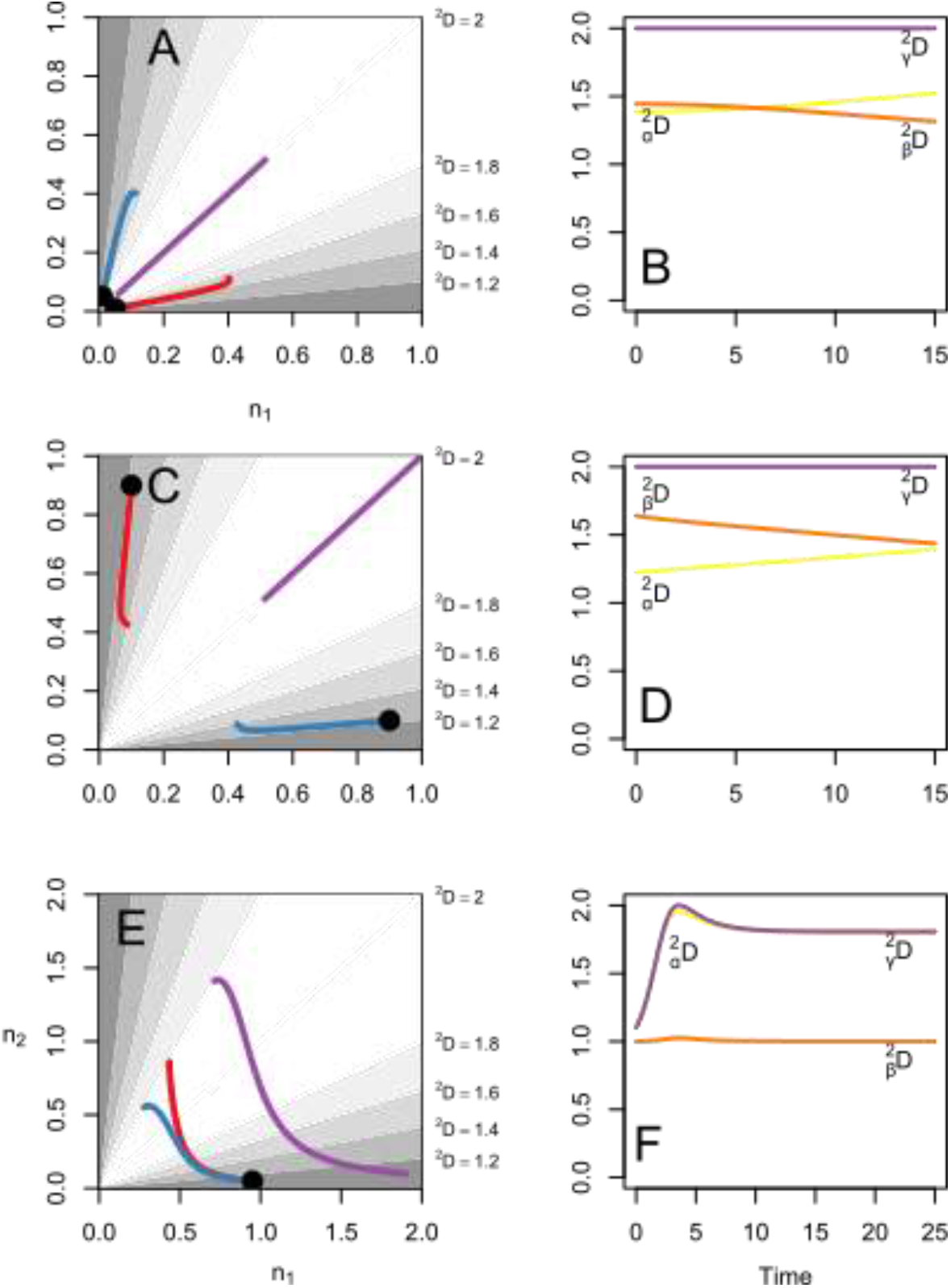
Illustrates how strong changes in absolute abundances can produce negligible changes in diversity across a metacommunity. A) Population growth in in a metacommunity (purple) consisting of two communities (red and blue) with two competing species. Shading denotes Simpsons’ diversity. Diversity changes little in either community or the metacommunity. B) Plotted over time, gamma diversity remains constant (purple), while alpha diversity increases slightly (yellow) at the expense of beta diversity (orange). C) Illustrates population decline in a metacommunity of two competing species. D) The resulting declines due to competition produce modest changes in alpha, beta or gamma diversity. E) A metacommunity where interactions are mutually beneficial in one community (red) and antagonistic in another community (blue). F) In spite of difference in interaction types, the changes in relative abundances run parallel across communities leading to no change in beta diversity. Alpha diversity and Gamma diversity follow roughly the same course and simulations within one community with mutualist (Main text Fig. 3 F) or one community with mutualists (Main text Fig. 3 H).

